# Calcium ions promote membrane fusion by forming negative-curvature inducing clusters on specific anionic lipids

**DOI:** 10.1101/2020.04.29.068221

**Authors:** Christoph Allolio, Daniel Harries

## Abstract

Vesicles enriched in certain negatively charged lipids, such as phosphatidylserine and PIP2, are known to undergo fusion in the presence of calcium ions without assistance from protein assemblies. Other lipids do not exhibit this propensity, even if they are negatively charged. Using our recently developed methodology, we extract elastic properties of a representative set of lipids. This allows us to trace the origin of lipid-calcium selectivity in membrane fusion to the formation of lipid clusters with long-range correlations that induce negative curvature on the membrane surface. Furthermore, the clusters generate lateral tension in the headgroup region at the membrane surface, concomitantly increasing its Gaussian bending modulus. Finally, calcium binding also reduces the orientational polarization of water around the membrane headgroups, potentially reducing the hydration force acting between membranes. Binding calcium only weakly increases membrane bending rigidity and tilt moduli, in agreement with experiments. We show how the combined effects of calcium binding to membranes lower the barriers along the fusion pathway that lead to the formation of the fusion stalk as well as the fusion pore.

## I. INTRODUCTION

Membrane fusion is a fundamental and vital biophysical process. Several well-known examples of fusion in cells are enabled by the SNARE machinery, which acts to empty the contents of vesicles from the cytosol into the extracellular matrix.[1, 2] The established crucial role of calcium in SNARE mediated fusion[3, 4] may be significant beyond its regulatory role in binding to the protein complex itself. Indeed, for certain membrane compositions, the calcium cation, Ca^2+^, is known to trigger membrane fusion on its own, i.e. even in the absence of any proteins.[5–7] The study of calcium-lipid interactions and their role in membrane fusion allows us to gain deeper insight into the physics of the fusion process, and to identify the underlying principles that may be obscured by the staggering complexity of the full protein assemblies. Moreover, identifying the driving forces of membrane fusion can also help make additional predictions about the action performed by the more complex cellular machinery.

In a recent study of Ca^2+^ and cell-penetrating peptide mediated membrane fusion by Allolio et al.[8], the entire fusion process was simulated in atomistic detail using DOPE/DOPS enriched membranes, starting from a high energy deformation of the bilayers in contact. It was shown that Ca^2+^ addition led to a destabilization of the membrane interface and subsequent exposure of the hydrocarbon tails to the solvent, thereby initiating stalk formation. While this nonequilibrium simulation provided unique phenomenological insight into the fusion process, as a virtual experiment it cannot provide the physical mechanistic insights to the process. In other words, on their own, simulations lack a fundamental molecular framework that can be generalized to other cases, such as additional lipids and proteins and moreover the free energies of the process are unknown. All simulations provide are the atomistic models used to generate the simulation and the final membrane remodeling outcome. Thus, in order to make predictions for other systems, it would be necessary to repeat the (computationally very expensive) simulation. Even if one could be satisfied by merely reporting such a large-scale simulation, the larger protein assemblies involved in mediating large-scale membrane deformations are beyond the scope of atomistic simulations, necessitating the use of a more coarsely-grained approach.[9]

Membrane remodeling processes have been very successfully described using the continuum Helfrich theory of curvature elasticity, and its later developments.[10–13] In particular the introduction of the lipid tilt degree of freedom by Hamm and Kozlov[14] has been effectively applied to membrane fusion, as tilt was found to be crucial in stabilizing the fusion stalk.[15] The Hamm-Kozlov-Helfrich (HKH) theory yields a free-energy functional with few, easily interpretable parameters and allows to evaluate the importance of each of the parameters for the fusion process by gauging its influence on the energetics of the process. As its application only requires the solution of a low dimensional variational problem, the HKH theory can be used to describe very large systems. The application of HKH theory to calcium mediated fusion therefore allows to address the large scales required to probe fusion events while at the same time furnishing an interpretation of the molecular structure in terms of continuum quantities.

In this vein, continuum theory has already shown fusion to be facilitated by properties such as lipid negative intrinsic curvature [6, 16, 17] and membrane tension[18, 19]. When a curvature elastic theory is applied to a membrane remodeling process, the effect of proteins is traditionally taken into account either as a scaffolding boundary condition[20], a rigid adsorbing surface[10, 21] or as an ideal insertion[22, 23]. The influence of interfacial interactions on the continuum parameters are generally neglected, mostly due to the lack of a systematic way to incorporate them into the theory. Yet, as a small ion Ca^2+^, is incapable of membrane remodeling by scaffolding or insertion. Thus, inasmuch as calcium influences membrane properties, it must do so through its high charge density as well by its strong binding to the membrane headgroups. In this study, we use molecular dynamics simulations to resolve how Ca^2+^ modifies membrane properties and show how in turn these modifications lower the vesicle fusion barrier in a lipid specific way, most notably by changing the spontaneous curvature of the membrane.

## II. RESULTS AND DISCUSSION

### A. Continuum Theory

The HKH free energy functional[14, 24] of membrane elasticity, relates the physical properties of lipid monolayers and flat bilayers to the energy of their deformations,

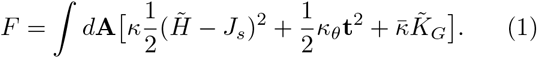

The energy density in this functional is determined by the bending modulus *κ* that is associated with the mean curvature 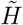 (note that the ^~^ signifies that the calculation is performed over the director field, rather than the membrane normal vectors); the tilt modulus, *κ_θ_* and the spontaneous curvature, *J_s_*. The Gaussian bending modulus 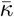 is usually irrelevant, because the integral of the Gaussian curvature 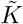 is a topological constant, according to the Gauss-Bonnet theorem. The HKH functional has been very successful in modeling membrane fusion, suggesting that knowing the pertinent elastic parameters should reveal the propensity of a given membrane towards fusion also when lipid properties are modified, for example in the presence of Ca^2+^. In particular, the effect of Ca^2+^ on membranes is known to be highly lipid specific and this should be reflected in the parameters.

To employ the HKH theory as a predictive tool, we have examined a representative set of membranes: neutral as well as anionic membranes, both formed of single components and containing mixtures, including phospholipid membranes with saturated tails (DMPC), hybrid lipids (POPC) and fully unsaturated tails (DOPC) as well as ceramides. Importantly, we studied the negatively charged phosphatidylserine (PS), as well as the negatively charged glycolipids PIP2 and GM1, all of which are known to interact strongly with Ca^2+^. We used a lipid:ion mole ratio of ≈ 1:10. Where more ions were required to ensure charge neutrality, we neutralized the system with Ca^2+^.

### B. Bending and Tilt

The influence of ions on membrane elasticity has been examined in a number of studies, with no current consensus. Poisson-Boltzmann and other mean field theories predict a weak increase in *κ* due to salt ions, which is nonlinear in salt concentration.[25–29] However, mean field theory is known to often fail for the interaction of polyvalent ions with interfaces, due to the rise of strong coupling between charges in the Coulomb fluid.[30, 31] Coarse grained simulations have found a decrease in the bending modulus for polyvalent ions, while confirming the weak increase for monovalent ions.[32] Experimental data on the other hand show an increase in membrane thickness through the addition of calcium[33, 34]. Because the bending modulus for a particular lipid is determined by the packing constraints on the chain in the lipid[35], any increase in membrane thickness generally corresponds to an increase in *κ*.[36] To the best of our knowledge the effect of divalent ions on the tilt modulus has not been studied at all. Our recently developed ReSIS methodology provides access to the membrane elasticity of charged lipid mixtures interacting with Ca^2+^ from atomistic simulation data.[37] In Fig. 1, upper panel we show the resulting bending moduli for all lipids examined in this study; monolayer thicknesses and areas per lipid can be found in the SI.

**FIG. 1.**
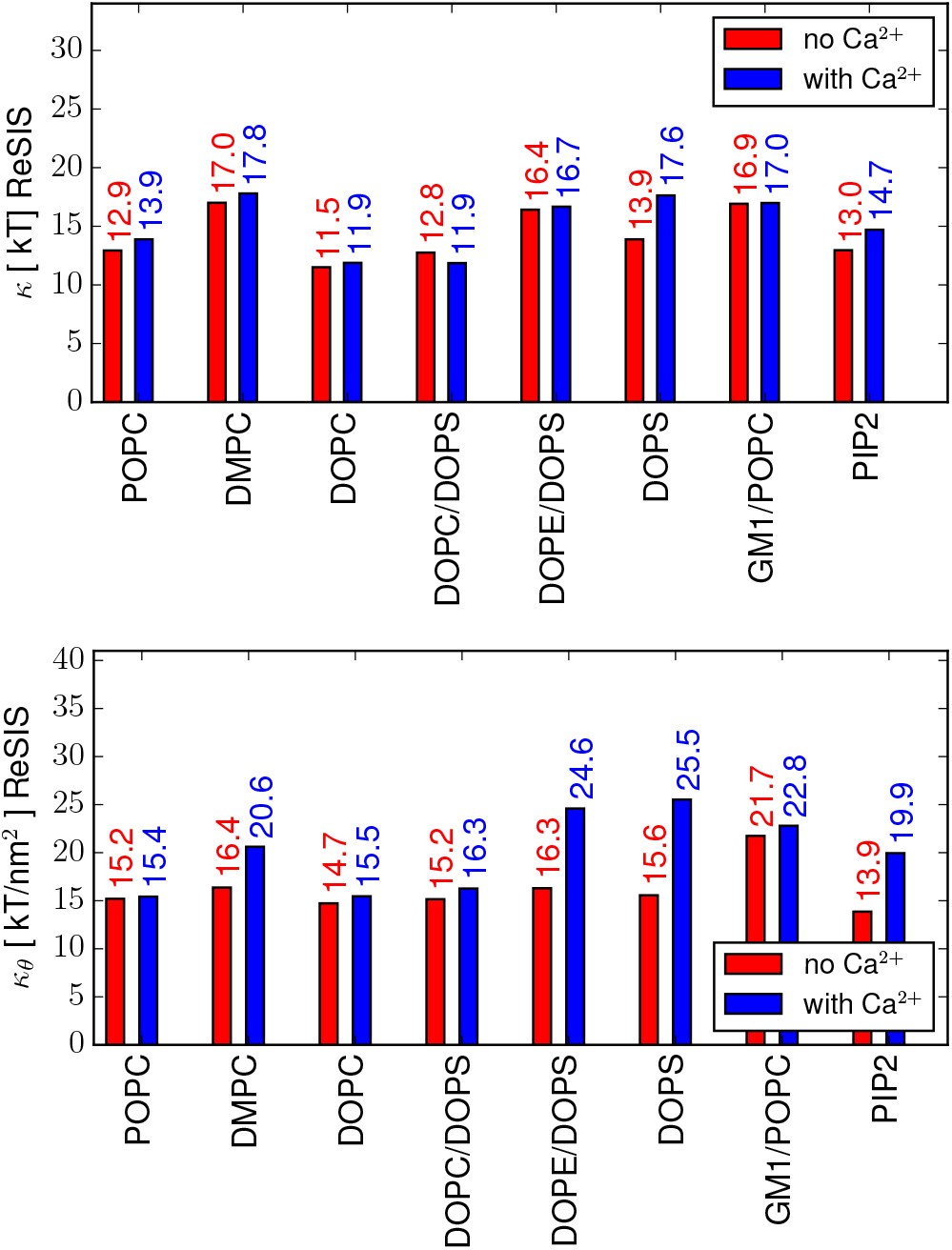
Elatic moduli for various lipids. Upper panel: Monolayer bending moduli of selected membranes in units of kT, in the absence of ions (or *K*^+^ counterions only, for charged lipids) and exposed to a CaCl_2_ solution with one Ca^2+^ ion per ten lipids, or full counterion charge in case of charged lipids. Lower panel: Tilt moduli for the corresponding membranes given in kT/nm^2^, reported per bilayer.

We find that Ca^2+^ barely increases *κ* in most membrane patches, with the only notable exceptions being pure PIP2 and PS lipids. Despite being charged, GM1 is among the least affected by Ca^2+^ binding. The tilt moduli shown in Fig. 1, lower panel surprisingly show a stronger trend than the bending moduli: in all membranes studied we observe an increase of the tilt moduli, exceeding the membrane stiffening observed in κ. The largest impact of calcium is found for DOPE/DOPS, DOPS, PIP2 and DMPC. This increase in tilt moduli should negatively affect fusion rates as stalk formation requires tilt[15], so the change in moduli is not an explanation for the increased propensity towards fusion observed in PS and PIP2. The surprising reaction of DMPC to the presence of Ca^2+^ in simulations is also reflected in the formation of gel-phase domains triggered by the addition of the ion at 303 K. While this is clearly an unphysical artifact due to overbinding by the Ca^2+^ ions in simulations, a lowering of the melting temperature of DMPC due to calcium has indeed been measured experimentally.[38] We have been able to reproduce gel phase formation using a charge-scaled force field, aimed to correct for some of the overbinding (see SI). Yet, it should be noted that in experiments too, Ca^2+^ binds more strongly to DMPC than to unsaturated lipids.[39] While the stiffening superficially confirms the mean field prediction for membranes in presence of salt, the origins of this effect must be very different: Because Ca^2+^ ions bind to the charged membranes, we should expect a softening as the charged headgroups are neutralized and counterion layers are removed. Instead, in our simulations we observe an increase in membrane rigidity, accompanied by membrane thickening.

### C. Monolayer Spontaneous Curvature

Once the monolayer values of *κ* are known, the first moment of the lateral pressure distribution *π*(*z*) allow to determine the monolayer spontaneous curvature *J_s_* by using the following relation.[14]

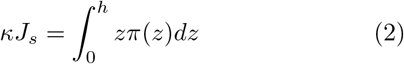

Here, the integration is carried out along the membrane normal from the center of the bilayer to the bulk. Thus, to determine the impact of calcium on monolayer spontaneous curvature, we first calculated the membrane lateral pressure profiles. In order to obtain the correct pressure contribution from the ionic double layer, we compute the stress tensor using particle mesh Ewald summation.[40, 41] This step is required to account for the extremely long range virial contributions of the charged ion layers (see Methods, and for further discussion see the SI).

Spontaneous monolayer curvatures obtained from Eq. 2 are given in Fig. 2, upper panel. In the absence of calcium, the spontaneous curvatures we calculated for the different membranes are close to experiment. Specifically, all PC membranes show low negative curvature[42], while GM1 shows positive curvature[43, 44]. Previously, PS has been measured to have positive spontaneous curvature, when deprotonated at pH 7.[45] This measurement was made using PE/PS mixtures in the hexagonal phase. In our PE/PS mixture, we find significant curvature overall, but slightly less negative than expected for pure DOPE (0.3-0.4 nm^-1^).[42, 45] This is in agreement with the experimental observation of increased spontaneous curvature in the presence of PS. Surprisingly however, the spontaneous curvatures of the pure anionic lipids PS and PIP2 are slightly negative. Especially at low concentration PIP2 is known to be stable in micellar aggregates in solutions, indicating positive *J_s_*.[46] A possible explanation for this apparent disparity is the confinement of the counterions to a small volume between lipid slabs due to the finite size simulation: The K^+^ counterions contribute to the lateral pressure (see also 2, lower panel) and have an effect on *J_s_* proportional to their distance to the membrane center *z*. Owing to their high concentration they also adsorb to a significant extent (see SI). At low concentration, these ions will be dissolved and reside far from the membrane, thereby increasing the first moment of the lateral pressure and hence the spontaneous curvature. Moreover, due to the finite size and limited hydration, lipids in simulations are expected to be under significant osmotic stress. Our results indicate, that structurally (i.e. in absence of net charge) PS and PIP2 lipids have negative curvature, which is overcompensated by electrostatics. Experimentally, it has also been shown, that at pH 2 (protonated) PS exhibits strongly negative *J_s_* in PE/PS mixtures, whereas in pH-neutral solutions, deprotonated PS has a very small *J_s_*.[45] There are indications that similar effects are also present in PIP2.[47]

**FIG. 2.**
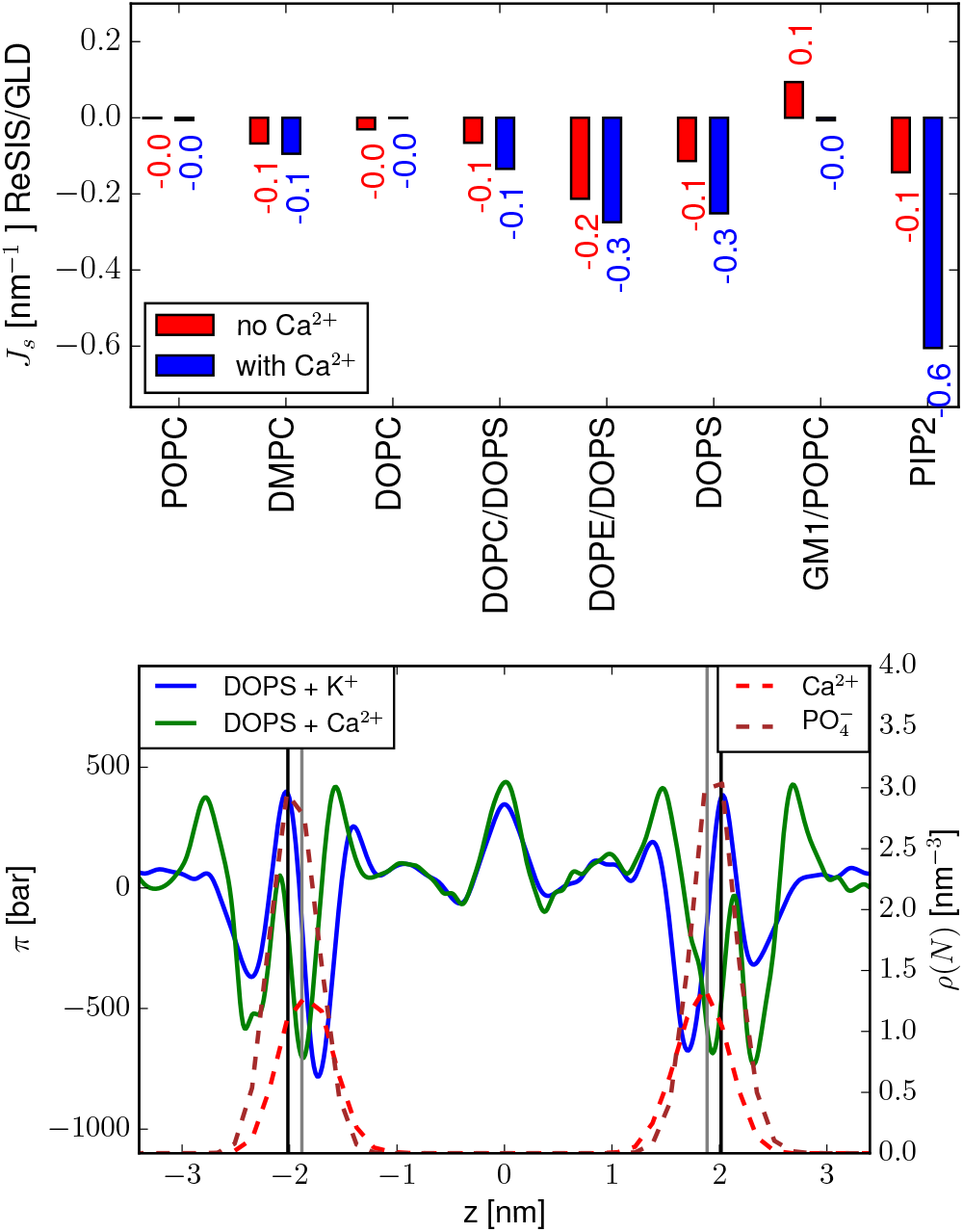
Impact of calcium on sponatneous curvature. Upper panel: Spontaneous curvatures for selected membranes, red bar no ions (or *K*^+^ counterions respectively, blue bar, exposed to a CaCl solution with one Ca^2+^ ion per ten lipids, or full counterion charge respectively. Lower panel: Stress profile of a DOPS bilayer in the presence of K^+^ vs. Ca^2+^ counterions (left axis) and number density profiles for Ca^2+^ ions and phosphorus atoms. The vertical lines represent the position of the water midplane (Gibbs-Luzzatti surface) of the membrane bilayers, in the presence of K^+^ (grey line) and Ca^2+^ (black line) ions.

Recent studies have debated whether Ca^2+^ induces positive or negative curvature on lipids.[48–51] In our simulations, we find that Ca^2+^ very strongly induces negative spontaneous curvature on negatively charged (pure) DOPS or PIP2 membranes. This is of great significance for fusion, as reduced spontaneous curvature is well known to lower the energy of the fusion stalk and pore.[11] Due to the finite size and ion confinement effects mentioned, the decrease of *J_s_* is likely to be even more striking at low ion concentrations.

In Fig. 2, lower panel we show the lateral pressure profile of DOPS in presence and absence of Ca^2+^. The pressure profile is strongly modified in the head group region from positive (in presence of K^+^) to negative pressure (in presence of Ca^2+^). The negative pressure is strong at the z-position where Ca^2+^ ions are concentrated and drives membrane thickening by Ca^2+^ as well as induces spontaneous curvature. As a rule, neutral membranes display a much smaller change in *J_s_* upon Ca^2+^ addition. The effect of Ca^2+^ on the curvature, of e.g. the negatively charged GM1 lipids is also small, despite full binding of the ions to the lipid (see Supporting information). Closer examination reveals that, the binding of Ca^2+^ to PS is complex, with significant contributions from the COO^−^, 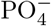 and carbonyl groups (see also Ref. [52] and the SI). It is therefore impossible to explain the molecular mechanism of curvature induction by using simple arguments based on charge densities alone. The overlap of peaks in the pressure profile with the density profiles of calcium ions and PO_4_ groups, as shown in Fig. 2 does not do justice to the full complexity of the molecular origins of the curvature induction, necessitating a more detailed investigation of headgroup binding.

### D. Role of Lipid-Calcium Clusters

We have already shown that the curvature of some lipid membranes is strongly affected by calcium, while others are almost unaffected. For some membranes, such as POPC, calcium binding is weak, which explains the small effect on curvature. Other PC lipids have stronger interactions (at least in simulation), but still exhibit a low influence of calcium on spontaneous curvature. Thus, the strength of binding to the lipid head groups clearly is not the only cause of curvature induction. Instead, we propose that long-range lipid clustering by calcium is responsible for the change in interfacial tension and the induction of negative curvature. To illustrate this, we define a local pair correlation function in the volume of a spherical cap *g_c_*, that allows us to selectively sample within the interfacial region (see the SI for details). The difference in the head group-head group correlation before and after Ca^2+^ addtion, Δ*_gp,p_* is a measure of induced headgroup ordering. The Fourier transform of this function ||Δ*_ĝp,p_*||, allows a visual estimate of the range of correlation induced by the ions. The bottom panel of Fig. 3 shows through the low *q* patterns modulation that the presence of Ca^2+^ induces long-range structure in the lipid head group region. This modulation is specific to lipids that exhibit negative curvature induction in the presence of Ca^2+^. This supports our hypothesis of the role of extended clusters.

**FIG. 3.**
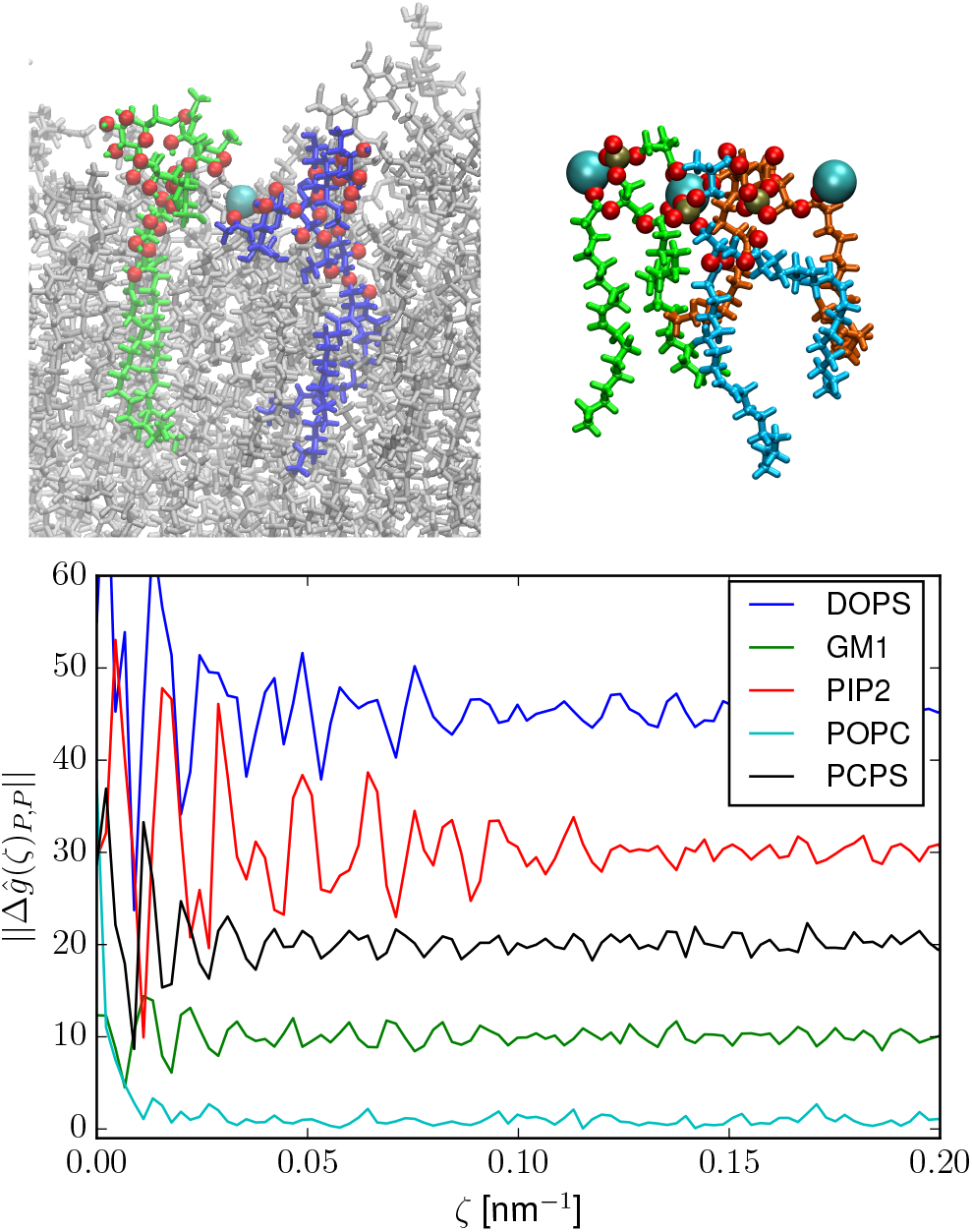
Top panel: Clustering of lipids by Ca^2+^ from simulations. Bottom panel: Correlation peaks obtained from the Fourier transformation of the difference of Interfacial phosphate-phosphate pair correlation functions. For legibility, the results for different membranes are shifted.

Why do some lipids form long-range clusters that strongly influence the mechanical properties of membranes while others do not? One particularly interesting case is GM1, which despite being negatively charged and shown to complex calcium in experiment[53] as well as in our simulation, hardly generates negative curvature nor affects the elasticity of the membranes. We find calcium to be bound to GM1 in two ways: either it is enveloped by the sialic acid sugar, or it clusters two GM1 molecules. Significantly, the negative charge of the sialic acid is positioned far from the membrane center, and is in contact with water through a well hydrated sugar moiety. (see Fig. 3, top right). This prevents the calcium ion from interacting with the inner-lying parts of the lipid bilayer, and presumably also prevents long-range mechanical coupling. This finding is confirmed by experiments, where GM1 lipids have been reported to have only slightly slower diffusion in presence of calcium[54] and are known to inhibit Ca^2+^-mediated membrane fusion[55]. In contrast, in simulations PIP2 binding to 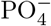 headgroups dominates. The Ca^2+^ ions form a network of bridges across the entire interface, which is facilitated by the exposure of multiple 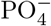 moieties of the lipid. Clustering of PIP2 by calcium is well established in experiment.[46]. PS also forms extended clusters (see Fig. 3, top left), as is also established experimentally[56]. For a full overview of ion binding see the SI.

### E. Barriers to Membrane Fusion

Having extracted from MD simulations all parameters that are used as inputs to the Helfrich elastic energy allows us to use these constants in the continuum theory and to evaluate the free energy of fusion stalk formation en route to fusion. With that, we can estimate the overall effect of Ca^2+^ on the energetic barrier for fusion stalk formation as well as identify the most important contributions to the energy barrier for fusion. Closely following the work of Kozlovsky et al.[15], we assume the stalk to have rotational symmetry and numerically solve for the minimal free energy surface, using finite tilt at the stalk origin and flat membrane surface at infinite distance as boundary conditions, see the SI for full details. The boundary condition of a high tilt at the stalk center is designed to model the fusing membrane interface. In Fig. 4, top panel the free energy of stalk formation is plotted as a function of *J_s_* for the different membrane compositions. The correlation with spontaneous curvature greatly overwhelms all other parameter changes. Fusion has a low barrier for the lipids which experimentally are found to undergo the fusion process, but prohibitively expensive (>30 kT) for those that do not. Specifically, we see that PIP2 forms stable stalks in presence of Ca^2+^ and that PS and DOPE/PS mixtures have a thermally accessible path to fusion. Taken together we have shown that stalk formation by Ca^2+^ ions can be successfully explained through its impact on curvature, and by extension provide a way to decode how fusion is triggered.

**FIG. 4.**
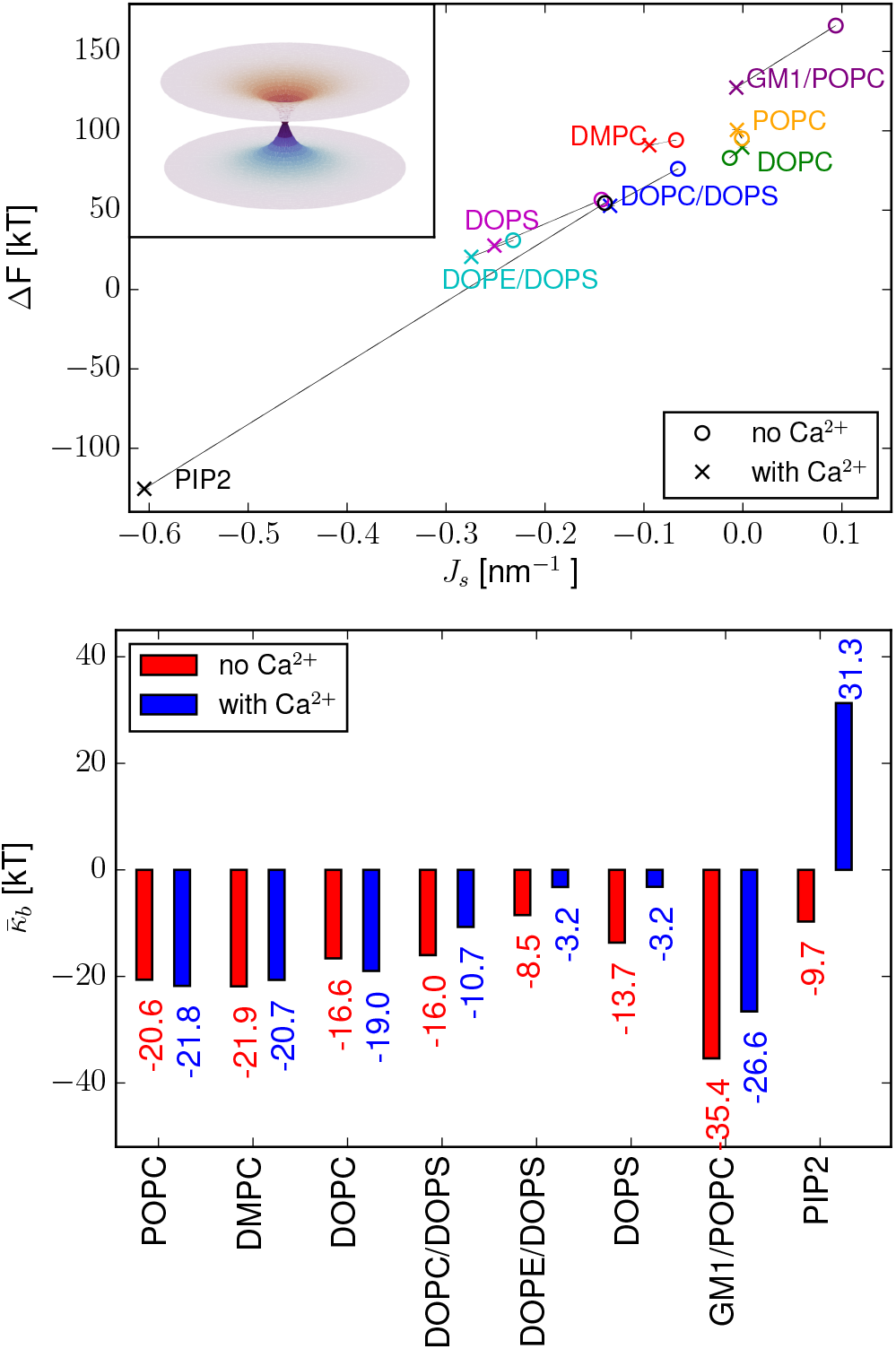
Top panel: Stalk formation free energies for different lipids at a stalk width of 40 nm, and stalk height of 2(d + 1). The stalk formation energy in the presence of Ca^2+^ is shown in blue, in the absence of Ca^2+^ it is given in red. Identical compositions are connected with grey lines. Inset: Example of stalk geometry. The stalk geometry does not depend strongly on the elastic parameters.[15] Bottom panel: Estimated values for the bilayer Gaussian bending modulus, from Eq. 5

However, stalk formation is only an early stage in the full fusion process: The fusion pore will cause a change in topology, which subsequently makes the Gaussian bending modulus contribution to the energy relevant. Importantly, the fusion pore can assume a catenoidal shape, that reduces the mean curvature to zero. This shape is expected, because with the opening of the fusion pore the bilayer shape of the membrane is restored. For symmetry reasons the spontaneous curvature of the lipid bilayer is zero, so that the Gaussian bending modulus becomes a dominant contribution in the energy density of the membrane.

A positive Gaussian bending modulus predicts stable phases with negative Gaussian curvature, such as cubic lipid phases or the stabilization of the opened fusion pore. The Gaussian bending modulus 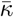 for the monolayers is very sensitive to increase in the lateral pressure *π*(*x*) in the headgroup region (even more than *κ*), as it is defined by:[14]

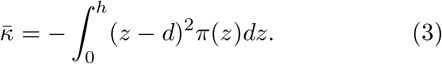

Furthermore, to evaluate 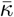, the position of the monolayer neutral plane at *d* is required. Unfortunately, eq. 3 does not seem to work well when used as a methodological basis for the extraction of Gaussian moduli from simulations. Specifically, the resulting monolayer Gaussian moduli are not in agreement with experiment for any physical values of d (see [57, 58] and references therein for in-depth discussion). To estimate the bilayer Gaussian bending modulus 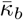, we instead begin with the typical experimental value of 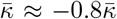 for the monolayer Gaussian bending modulus. The experimental ratio is conserved across lipids within 15%. For a membrane with neutral plane at distance d from the bilayer center any principal curvature of the monolayer *c_m_* is related to the principal curvature of the bilayer *c_b_* by

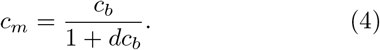

To first order in *d*[58], the bilayer Gaussian modulus is then given by

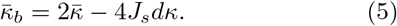

Therefore, by creating negative spontaneous curvature the Ca^2+^ ion can also significantly increase the Gaussian bending modulus. We computed 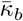 from Eq. 5, by inserting the values we obtained for *κ_m_*, d (the position of the neutral plane according to ReSIS lipid centers[37]) and *J_s_*. Alternatively, 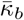 can be directly calculated by integrating the lateral pressure’s second moment over the entire bilayer (see SI). Both methods show that the sign of the Gaussian bending modulus can be inverted by Ca^2+^ for the PIP2 membranes (see Fig. 4, bottom). The pressure method also predicts an equivalent sign change for PS and a positive 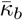 for the DOPE/DOPS system even without calcium (the latter further increases upon addition of Ca^2+^). This underscores the challenges of computing Gaussian moduli from pressure profiles. Contributions from thickening and stiffening also play a role, especially since clustering also thickens the membrane. Our robust conclusion, however, is that the generation of negative curvature by calcium mediated clustering also supports fusion by lowering the energetic cost of the topological change necessary for the formation of the fusion pore (unambigously turning into stabilization of the pore in the case of PIP2).

### F. Hydration and Electrostatics

So far we have determined the propensity for fusion in terms of negative curvature induced by clusters. However, from the perspective of kinetics, connecting two apposed bilayers through the transfer of a lipid tail into the adjacent interface is widely accepted as the rate determining step for stalk formation.[8, 59] For this lipid tail flip-flop to occur it is necessary to expose parts of the hydrophobic interface. This degradation of the interface is incompatible with the description of Helfrich theory, which assumes an intact membrane surface (interpretable as a thermodynamic average). The creation of a hydrophobic interface requires moving aside the far better solvated head groups. In a tensionless membrane, the tension in the headgroup region has to be counterbalanced by the tension of the chains

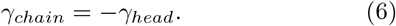

Disruption of the interface can be accomplished by creating tension on the membrane or in the headgroup region. Adsorption of Ca^2+^ also thickens the membrane, due to the contraction of the headgroups, as expected from the lateral pressure profile in Fig. 2. If the membrane cannot respond fully by lateral contraction (for example because it is a vesicle with internal pressure) a net tension results, which destabilizes the headgroup-water interface.

When membranes are in close proximity to each other, the need for a partial desolvation of their interfaces generates repulsive hydration forces, expected to dominate at very low intermembrane distances. Solvation of the interface by water molecules generates an orientational polarization of these water molecules. Before membranes can touch, this polarization has to vanish for symmetry reasons. Following Kanduč et. al.[60, 61] we therefore use the water orientation along the membrane normal as a proxy for the resulting hydration repulsion. In neutral bilayers only the water molecules very close to the dipolar headgroups (up to ca. 0.5 nm from the interface) show orientation along the membrane normal. The loss of orientation then closely tracks the decay of the hydration force.

In the top panel of Fig. 5, we show the orientation of water dipoles along the membrane normal in the DOPS system. Water responds to the presence of charged lipids with a net orientation in the bulk water phase. In this case, water orientation also contains information about the electrostatic repulsion between membranes. Indeed, we find that long-range water orientation is entirely due to surface charge. This is shown by the agreement of the orientation with the value predicted for a water dipole in the local electric field of the simulation box. Surface charge can be present either in the form of bound ions, or in the form of charged lipids in the absence of absorbed counterions. To account for the long-range effect of surface charge, we evaluate the integrated polarization of water. The result is shown in the bottom panel of Fig. 5. The effect of the electric field is clearly seen, as neutralizing the charged membranes with Ca^2+^ removes a large amount of water polarization. As expected, in neutral membranes, adsorbing Ca^2+^ produces positively charged interfaces and thereby generates long range polarization. Negatively charged membranes, on the other hand, can lose their mutual electrostatic repulsion when calcium is added. Strikingly, once neutralized by calcium, the fusable membranes also have a low water polarization when compared with regular charge-neutral membranes. We conclude that the formation of clusters also disrupts the dipolar solvation of the membranes, which coincides with the observed increase of membrane interface tension. Together with the tendency of calcium to cross-link across membrane leaflets this effect is expected to further facilitate stalk formation via a decrease in hydration repulsion. The fact that in presence of Ca^2+^ the negatively charged membranes become effectively charge neutral also justifies our neglect of electrostatic forces in the continuum description of membrane fusion.

**FIG. 5.**
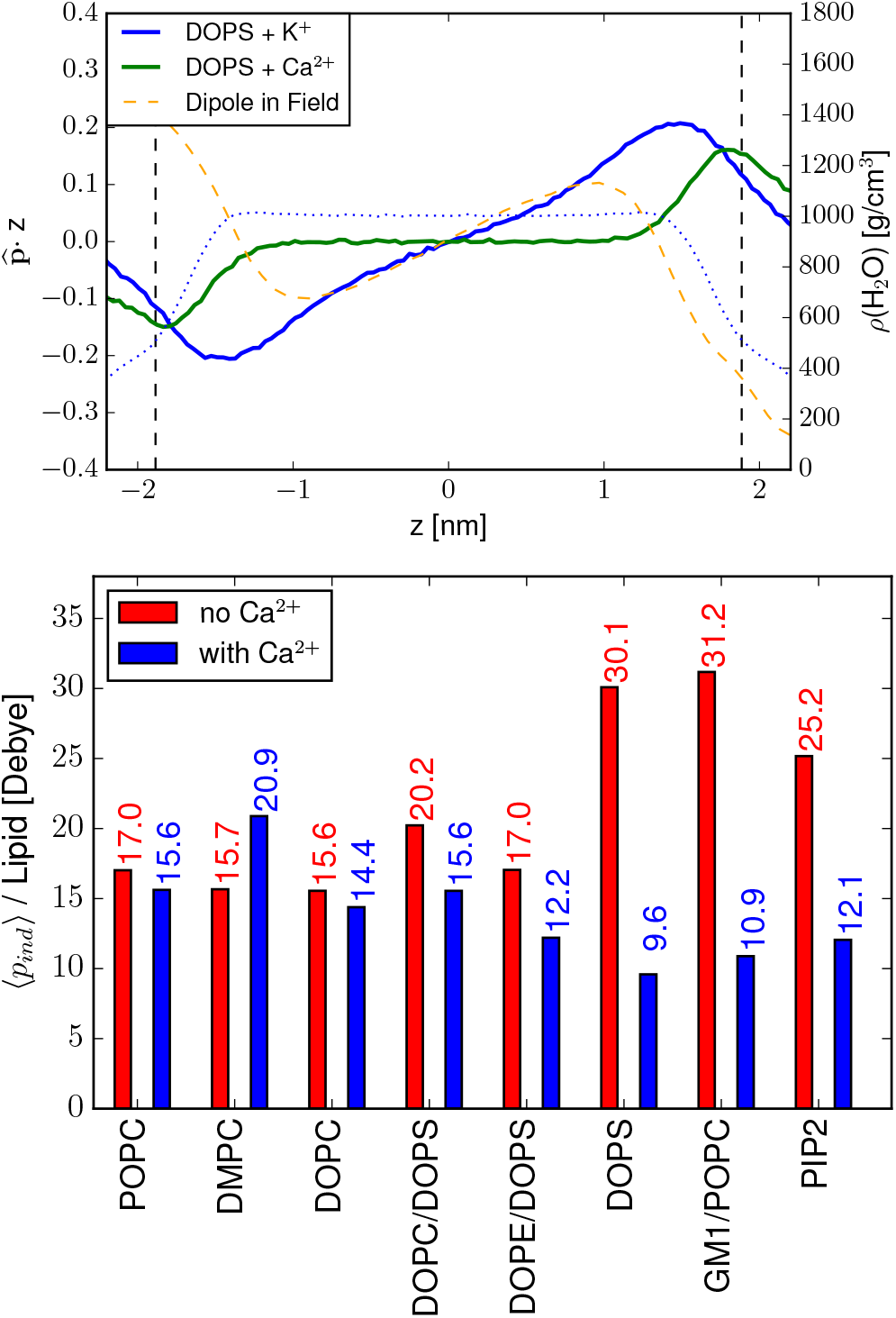
Top panel: Water orientation along the membrane normal, and computed dipole orientation generated from a Boltzmann distribution of the water dipole in the electrostatic potential of the Ca^2+^-free box. Bottom panel: Total polarization obtained by integrating over the orientation profile starting from the point where *ρh_2_o* = 350 mg/cm^3^.

## III. CONCLUSIONS

Using molecular dynamics, we have examined the mechanism of Ca^2+^ mediated membrane fusion in detail. We showed that, for certain lipids, Ca^2+^ selectively coagulates the membrane headgroups into long-range clusters in a lipid-specific way. Extending our recently developed methodology, we then extracted elastic parameters as well as the local pressure from molecular dynamics simulations. Thereby, we have been able to show how lipid-Ca^2+^ clusters generate membrane surface tension in the headgroup region, which in turn translates to a negative spontaneous curvature, an increased Gaussian bending modulus, and a depolarization of the water interface. Using continuum modeling, we then estimated fusion stalk formation energies and obtained the experimentally observed lipid selectivity. In the absence of net charge on the membrane interface, the induction of negative curvature by calcium is coupled to tension in the headgroup region, an increase in the Gaussian bending modulus, and reduced hydration. Curvature induction by adsorbates is therefore an easily available predictor for fusion and phase stability. The methods employed in this study are fully local and can easily be extended to membrane proteins, such as the SNARE complex, their surrounding lipids and other biophysical systems of relevance. By conducting accurate membrane simulations of modest size, and extracting continuum properties, we provide a new mode of access to large-scale deformations, and to probing the molecular details of membrane remodeling processes.

## IV. METHODS

All simulations were performed using the CHARMM36[62–64] all atom force field for the lipids, the TIP3P[65] model for water and semi-isotropic pressure coupling. We used the Roux calcium ions[66] with NBFIX corrections[67] as well as the scaled ECCR calcium[68] ions for comparison. Lateral pressure calculations use a modified version of the Sega code[40], that incorporates a Goetz-Lipowsky[69] force decomposition. Membrane shape optimization used a spline-based finite element approach. For full details on simulation setups, ion binding, surface variation, and pressure tensor evaluation see the Supporting Information.

## ACKNOWLEDGMENTS

CA thanks the Minerva foundation for a Postdoctoral Fellowship and the Charles University for a PRIMUS grant (PRIMUS/20/SCI/015).

