## Supporting Information for "Calcium ions promote membrane fusion by forming negative-curvature inducing clusters on specific anionic lipids"

### I. CONTINUUM MODEL

Our calculation of the stalk model follows the work of Kozlovsky and Kozlov.[1] We briefly outline and explain the different implementation based on their work. The ansatz assumes rotational symmetry of the stalk around the $z$ axis, i.e. the height. The stalk surface can be expressed as a function of the stalk radius $r$ and the angle $\psi$, which is the angle between the local tangent of the stalk surface and $r$. In the Hamm-Kozlov picture of Helfrich theory[2], the chain director is no longer necessarily the membrane normal, so that a tilt angle $\phi$ is required to recover the chain director $\mathbf{n}$. The divergence of this director is then used to compute the mean curvature:

$$\tilde{H} = \nabla \cdot \mathbf{n} = \cos\phi \frac{1}{r}\frac{\mathrm{d}(r\sin\psi)}{\mathrm{d}r} + \cos\psi \frac{1}{r}\frac{\mathrm{d}(r\sin\phi)}{\mathrm{d}r}. \tag{1}$$

Inserting this expression into the Kozlov-Hamm functional for the monolayer yields

$$f_m = \frac{1}{2}\kappa\left[\cos\phi\frac{1}{r}\frac{\mathrm{d}(r\sin\psi)}{\mathrm{d}r} + \cos\psi\frac{1}{r}\frac{\mathrm{d}(r\sin\phi)}{\mathrm{d}r} - J_s\right]^2 + \frac{1}{2}\kappa_t(\tan\phi)^2 - \frac{1}{2}\kappa J_s^2. \tag{2}$$

The spontaneous curvature $J_s$ and the moduli $\kappa_t$ and $\kappa$ have the same definitions as given in the main text. The final added term serves to establish a reference to the flat bilayer. The membrane bilayer consists of two monolayers which are part of the same membrane. Their geometry can be computed from the midplane of this membrane and the radius $r_c$. The *distal* monolayer $d$ is located away from the stalk, whereas the *peripheral* monolayer is $p$ placed at the point where monolayers touch. Following the normal vector at the midplane their radius is given as:

$$r_d = r_c - \delta\sin\psi_c; r_p = r_c + \delta\sin\psi_c \tag{3}$$

Where $\psi_c$ is the equivalent of $\psi$ for the bilayer midplane and $\delta$ is the monolayer thickness. Accordingly, the monolayers need to fulfill

$$\psi_c(r_c) = -\psi_d(r_d) = \psi_p(r_p), \tag{4}$$

which generates opposing curvature for each of the bilayers. The tilt functions $\phi$ remain independent for each leaflet. For the total energy, both monolayer energies have to be added and integrated over their respective surfaces. The resulting total free energy is

$$F = 2\pi\int\frac{\mathrm{d}r_c}{\cos\psi_c}\left[r_p\frac{\mathrm{d}r_p}{\mathrm{d}r_c}f_p(\phi_p,\psi_c) + r_d\frac{\mathrm{d}r_d}{\mathrm{d}r_c}f_d(\phi_d,\psi_c)\right], \tag{5}$$

where we have carried out a substitution of $r_{d,p}$ with $r_c$ and $\psi_{d,p}$ with $\psi_c$. This energy functional is easily recast as a Lagrangian.

$$F = \int \mathrm{d}r_c\, L(\phi_d, \phi_d', \phi_p, \phi_p', \psi_c, \psi_c', r_c) \tag{6}$$

The solution for the optimal surface can be obtained by the variational method, so that

$$\delta F \stackrel{!}{=} 0. \tag{7}$$

In order to simplify expressions, we follow Kozlov and Kozlovsky and substitute

$$z = \tan\psi_c; x, y = \sin\phi_{d,p} \tag{8}$$

The final stalk height is

$$H = \int \mathrm{d}r\, z(r). \tag{9}$$

This value can be constrained using an appropriate Lagrange multiplier. Unfortunately, the corresponding Euler-Lagrange equations are highly nonlinear and do not easily lend themselves to numerical solution.[3] Therefore, instead of solving the Euler-Lagrange equations, we constructed a numerical variational procedure, starting from an initial guess that fulfills the boundary conditions. The boundary conditions for $r_c \to \infty$ are vanshing tilt, splay and curvature: $\psi_c \to 0$, $\phi_{d,p} \to 0$. We begin by constructing the trial functions

$$\psi_c^i(r) = \psi_c^0 \exp\left[\frac{-\{(r-r_0)\sqrt{\pi}\psi_c^0\}^2}{H^2}\right] \tag{10}$$

and

$$\phi_{p,d}^i(r) = \phi_{p,d}^0 \exp[-\lambda(r-r_0)] \tag{11}$$

which fulfill the boundary conditions as well as the height constraint. In order to ensure a rapid decay of the tilt deformation, we set $\lambda = 3$. The initial conditions we chose as $\psi_c^0 = \pi/5$ and $\phi_{d,p}^0 = \pm\pi/5$. Because the radius of the distal monolayer $r_d$ must not be smaller than 0, we can set the lower integration boundary at

$$r_0 = \delta \sin \psi_c^0. \tag{12}$$

In addition, we also choose an upper integration boundary, determined as the stalk width. For this quantity we set an arbitrary value of 40 nm. In the next step, we use the ansatz

$$F = \int \mathrm{d}r_c\, L(\phi_d + \xi_d, \phi_d' + \xi_d', \phi_p + \xi_p, \phi_p' + \xi_p', \psi_c + \eta_c, \psi_c' + \eta_d, r_c). \tag{13}$$

Following Kozlovsky et al.[3], we construct the trial functions $\xi_{d,p}, \eta_c$ from base splines. In order to continue to fulfill the boundary conditions, these functions need to be zero at $r_0$. The base splines require a set of inner knot points, for which we chose equidistant spacing between $r_0$ and $R$. We used eleven inner knots. From these we constructed a cubic spline basis, so that e.g.

$$\xi_d(r) = \sum_{i=2}^{11} c_i B_i(r). \tag{14}$$

The first polynomial was removed to fulfill the boundary condition $\xi_d(r_0) = 0$. In order to ensure continuous differentiability of the resulting sum, a spline polynomial with extended boundaries was fitted on the sum of initial guess and trial function before differentiation. The integration was carried out using the standard quad method of SCIPY.[4] Having thus turned the variational problem into a multidimensional optimization problem, we solve for

$$\nabla F\{c_i\} = 0 \tag{15}$$

using standard techniques, such as conjugate gradients and quasi-Newton procedures as implemented in SCIPY[4]. During the optimization procedure we add the term

$$\lambda\left(\int \mathrm{d}r\, z(r) - H\right)^2 \tag{16}$$

with $\lambda = 5000$. After minimization, we remove the penalty function to give the final energy value. A typical result is given in Fig. 1. We observe an overshooting of the stalk height, the so called “wings” of the stalk in the case of a large integration boundary in agreement with what was observed in [1]. These wings can be eliminated by adding hydration repulsion at low stalk heights[3].

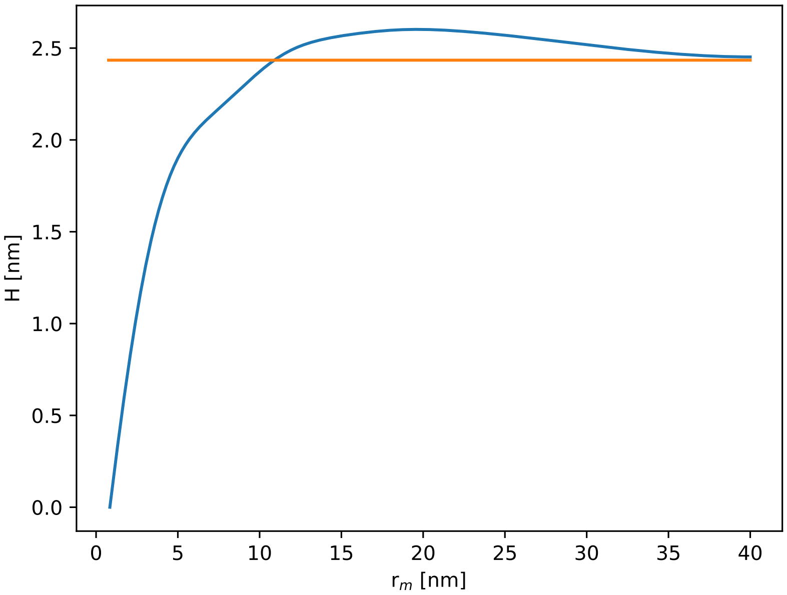


FIG. 1. Stalk of DOPS with adsorbed $Ca^{2+}$. The stalk geometry is known to depend very little on the membrane composition.[3] The stalk height $H$ is marked with the yellow line. We plotted the center between the two stalk monolayers.

### II. STRESS DECOMPOSITION

The stress tensor can be decomposed into a part that depends on the potential $\sigma_V$ and a part depending on the momenta $\sigma_K$, the kinetic stress tensor:

$$\sigma = \sigma_V + \sigma_K \tag{17}$$

The kinetic part of the stress tensor is available from the velocities, Irving and Kirkwood[5] have shown the local, molecular version of the kinetic stress tensor to be:

$$\sigma_K(\mathbf{x}) = -\left\langle \sum_i \delta(\mathbf{r}_i - \mathbf{x}) m_i \mathbf{v}_i \otimes \mathbf{v}_i \right\rangle \tag{18}$$

where the delta function is used to account for the locality of the atoms at positions $\mathbf{r_i}$, and the atomic velocities are given as $\mathbf{v_i}$. The brackets indicate thermal averaging, following Kreuzer.[6] The potential part of the stress tensor requires the integration of the force between atoms along a contour $\mathcal{C}$, such that its components (indiced by greek letters) are given by

$$\sigma_V(\mathbf{x})^{\alpha\beta} = \frac{1}{2}\left\langle \sum_{i\neq j} f_{ij}^{\alpha} \int_{\mathcal{C}_{i,j}} \delta(\mathbf{x} - \mathbf{l}) ds^{\beta} \right\rangle. \tag{19}$$

The equation is given here in terms of pairwise forces $f_{ij}$ and the integration is carried out along a contour connecting the two atoms $i$ and $j$ with line element $\mathbf{l}$. The choice of contour is arbitrary, which appears as a consequence of the stress tensor being determined only up to a divergence-free field.[7] Unfortunately, even though the contour is an arbitrary parameter, in the general case the results of the theory depend on it. Typically the Irving-Kirkwood-Noll[5, 8] contour is chosen, i.e. a straight line between the atoms. In the Harasima contour[9], the integration is carried our from $\mathbf{r_i}$ first along a straight line in the $xy$ plane to the $xy$ position of $\mathbf{r_j}$ and then along the $z$ axis to reach $\mathbf{r_j}$. We then discretize the result to a set of slices along the $z$ direction (see for example Ref. [10]). The resulting averaged expression, for the virial $\sigma_V$ corresponding to the lateral stress $x$ and $y$ is:

$$\sigma_L(z) = \frac{1}{2V_s}\left\langle \sum_{i\neq j} \frac{f_{ij}^x r_{ij}^x + f_{ij}^y r_{ij}^y}{2} \Theta(z_{su} - r_i^z)\Theta(r_i^z - z_{sl}) \right\rangle \tag{20}$$

where $\Theta$ is a Heaviside step function and $z_s$ represent the boundaries of the slab. In the summation half of the lateral stress contribution is assigned to the slab on each atom. This formulation is particularly suited for reciprocal space, as the instantaneous contribution to the stress of the particle $i$ by Ewald summation may be expressed as[10]:

$$\sigma_{K,i}^{\alpha\beta} = -\frac{q_i}{V^2\epsilon_0}\sum_{\mathbf{K}\neq\mathbf{0}} Q(k)\,\mathrm{Re}[\exp(-i\mathbf{k}_n\cdot\mathbf{r}_i)S(\mathbf{k}_n)]\left[\delta_{\alpha\beta} - 2k_n^{\alpha}k_n\beta\left(\frac{1}{k_n^2}+\frac{1}{4\kappa^2}\right)\right] \tag{21}$$

Naturally, the binning and thermal averaging have to be performed just as in Eq. 20. Here $\mathbf{k}$ are reciprocal lattice vectors, $\kappa$ is the shift factor between real space and reciprocal space and $\delta$ here is a Kronecker-delta instead of the Dirac distribution used everywhere else in the manuscript. The functions $S$ and $Q$ are defined as follows:

$$S(\mathbf{k}_n) = \sum_{j=1}^{N} q_j \exp(i\mathbf{k}_n\cdot\mathbf{r}_j) \tag{22}$$

$$Q(k_n) = \exp\bigl(-k_n^2/4\kappa^2\bigr)/k_n^2 \tag{23}$$

This approach can be incorporated to a formulation of PME (particle-mesh Ewald summation), as was implemented and validated by Sega.[11] We use Sega's implementation of the PME Harasima stress. Coulomb and Lennard-Jones potentials are two body functions, so the the forces $\mathbf{f}_{ij}$ are easily obtained. Force fields do contain many-body potentials as well, for these we implemented a Goetz-Lipowsky decomposition (GLD).[12] This decomposition was chosen, because it implies a degree of averaging over contours and is numerically stable for any potential. Its form is

$$\mathbf{f}_{ij} = \sum_{\langle j\rangle}\frac{1}{n_M}\left(\frac{\partial V_M}{\partial\mathbf{r}_i} - \frac{\partial V_M}{\partial\mathbf{r}_j}\right). \tag{24}$$

Here $n_M$ is the number of bodies in the potential, and the summation is carried over all the pairs of particles in the potential. In contrast to the GLD, the central-force decomposition (CFD) has the favorable property of generating symmetric contribitions to the stress tensor. However, the CFD's behavior is ill defined in situations of linear dependence of force vectors in many-body potentials. Especially, for dihedral potentials at flat angles, which do not correspond to extrema in the potential.[13] These occur in the CHARMM36 lipid force field, so that this force decomposition could not be used. As a side remark, the symmetry of the stress tensor itself is controversial for molecular systems, even more so in the presence of many-body potentials. Going into depth here is beyond even the scope of this SI, so the reader is referred to the literature.[7, 13, 14] In practice, the Harasima contour has the additional drawback of not producing useful normal pressures. In order to verify its usability for our case we compare it to results with an IKN contour in the case of an uncharged membrane, as shown in Fig. 2. For heavily charged membranes we are inclined to trust the effect of PME over the effect of the contour. The good agreement between Harasima and IKN lateral pressures for slab systems has often been documented in the literature, so we feel justified in making this choice.[10, 11, 15] The normal pressure has to be uniform, for reasons of mechanical stability. As our simulations were run at zero tension, we impose

$$\int dz[p_L(z) - p_N] = 0. \tag{25}$$

This choice was validated by comparing the normal pressure thus obtained with the one from the virial, yielding in general only a small difference (0-30 bar). In addition, we had to deal with lack of an implementation of the SETTLES[16] algorithm in the Sega code. It was later discovered, that the SETTLES are also a significant source of normal pressure imbalance in the current reference IKN implementation. Therefore, for the pressure calculation, we substituted the SETTLES with triangular water molecules, constrained by LINCS[17]. In order to achieve reasonable results, the LINCS order had to be increased to 5 in the rerun. The resulting force profiles were then spline-interpolated and the momentum integration was taken between bulk and membrane center. We averaged over 20000 frames taken from a rerun of the last within 200 ns of sampling obtained from standard version of GROMACS 5[18], imbalances in the pressure profile arise from the asymmetric adsorption of $Ca^{2+}$. We took the average of the moments calculated on both leaflets.

### III. INTERFACIAL RDF

#### A. Derivation

We define an interfacial radial distribution function (RDF), with the aim of sampling interfacial structure. Ordinary RDFs are widely used and can be easily interpreted by scientists familiar with structure factors. To sample the

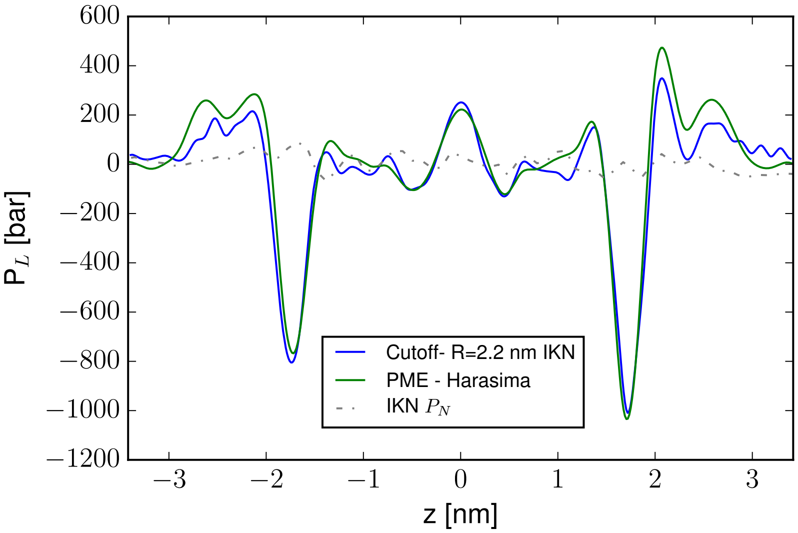


FIG. 2. Comparison of pressure profile using IKN and Harasima contours. The Coulomb cutoff for the IKN procedure is 2.2 nm. The IKN normal pressure is plotted. The moments of the pressure distribution are -1.82 kT/nm (Harasima) vs -2.28 kT/nm (IKN) respectively. The data is from a separate run without NBFIX.

structure within an interface it is practical to restrict the sampling to the particles inside the interface (see Fig. 3). In particular, 2D RDFs (e.g. with the radius containing only x and y components) are contaminated with information from the bulk and from the opposing leaflet, as are ordinary RDFs of bound ions.

Following Ben-Naim[19] we begin with the generic pair distribution function

$$\rho^{(2)}_{AB}(\mathbf{X}, \mathbf{X}')d\mathbf{X}d\mathbf{X}' \tag{26}$$

which describes the probability to find one particle of type A in the section of space $d\mathbf{X}$ and a particle of type B in the section $d\mathbf{X}'$. In particular we define the pair-correlation function $g_{AB}$ by

$$\rho^{(2)}_{AB}(\mathbf{X}, \mathbf{X}') = \rho^{(1)}_A(\mathbf{X})\rho^{(1)}_B(\mathbf{X}')g_{AB}(\mathbf{X}, \mathbf{X}') \tag{27}$$

Assuming the sytem is fluid, we expect that for long distances $g_{AB}(\mathbf{X}, \mathbf{X}') \to 1$, which means the density of particle $B$ at $\mathbf{X}'$ is uncorrelated to the density of particle $A$ at $\mathbf{X}$. If the fluid (interface) is also homogeneous, then it is possible to set an arbitrary origin at $\mathbf{X}$ and then write $g_{AB}(R(\mathbf{X}, \mathbf{X}'))$, so as a function of only the distance between the particles. Otherwise this is not generally possible, but one loses some information when transforming to

$$\rho^{(2)}_{AB}(R) = \rho_B g_{AB}(R) \tag{28}$$

In the next step, we assume that our system consists only of a particular region $\mathbf{X}, \mathbf{X}' \in S$ in which the interface is located. This region corresponds to an open system, so that the probability is replaced by an average number of particles. In particular, if the bulk is large enough, the interfacial system is in a grand-canonical ensemble, with a chemical potential of particles inside the interface equal to that of the corresponding particles in the bulk. Obviously we can still define a $g'_{AB}(\mathbf{X}, \mathbf{X}')$, which is arbitrary outside the region, but inside will be equivalent to the definition for the full system. Also this is still reducible to $g'_{AB}(R(\mathbf{X}, \mathbf{X}'))$, for the points in the region provided that the matter inside the region is a homogeneous fluid. This condition is not generally fulfilled for fluid interfaces. Nevertheless, e.g. for a flat interface along $z$ translational symmetries exist in the $x, y$ directions and rotational symmetry exists in $\phi$, so the expression should become exact for small enough intervals $dz$ and $d\theta$ respectively. In particular, in an (open) bulk system $g'_{AB}(R)$ is the same as in the bulk over the entire domain. The number of particles of type B in a spherical volume $dR$ around particle $A$ at $\mathbf{X} = 0$ is

$$N_{CN}(R') = \rho_B \int_0^{2\pi} \int_0^{\pi} \int_0^{R'} g_{AB}(R)R^2 dR d\theta d\phi \tag{29}$$

which is known as the coordination number. Note, that the integration order can be chosen at will, provided the limits are chosen correctly, however only the innermost integral limit can depend on a function, so that the outermost limit has to go over all possible radii. For the region S, we chose to carry out the integration in R last:

$$N'_{CN}(R') = \rho_B \int_S \int \int g'_{AB}(R) R'^2 d\theta d\phi dR \tag{30}$$

The coordination number $N'_{CN}$ includes only atoms in the interfacial region $S$. As $g'_{AB}(R)$ depends only on $R$, regardless of the actual form of the limits, it will be integrated as a constant over these two coordinates. Before the final step this part will therefore accumulate a prefactor $\frac{dV_S(R)}{dR}$, with $V$ being the Volume of region S. We include the Jacobian contribution $R^2$ into this volume, because this removes coordinate system dependency. The final integration will therefore be:

$$N'_{CN}(R') = \rho_B \int_0^{R'} \frac{dV_S(R)}{dR} g'_{AB}(R) dR \tag{31}$$

The coordination number can be extracted from simulation by simple counting of adjacent particles in slices, up to to the cutoff $R'$. Up to this point the discussion has been completely general to any region, provided it is sufficiently homogenous. As we only have to define $V(R')$, we chose the geometry in Fig. 3, where the interface consists of a flat region, limited by water on the outside (high $z$ and lipid on the inside low $z$). The volume of a spherical cap is known[20] to be

$$V_C = \frac{\pi h^2}{3}(3r - h), \tag{32}$$

with $h$ as the height of the spherical cap. This volume has to be subtracted from the full volume of the sphere, so that

$$\frac{dV_S(r)}{dr} = 4\pi r^2 - \pi h^2 - \pi h'^2 \tag{33}$$

where the subtraction only occurs if $r > h$ or $r > h'$ respectively.

$$h = p_z + r + d - Z; h' = r + d' - p_z \tag{34}$$

With $d, d'$ defined as distances along the $z$ axis from the box limits as shown in Fig. 3. $p_z$ is the $z$ position of the particle and $Z$ the extension of the box in $Z$ direction. The origin of the $z$ axis was set to the membrane center. This center was computed using the $z$ positions of the membrane lipids as computed from the lipid definitions used for the ReSIS surfaces[21]. In this way sampling was reduced to one half space along the local membrane normal, taking the membrane center as the origin. The interval defining the interface was $d, d' = 1\,\text{nm}$ In other words, the pair distribution function includes roughly only the interfacial region, as there is ≈2 nm of bulk water density, and a membrane thickness of ≈3 nm. We use these cutoffs to compute $g'_{AB}(R)$ from the coordination numbers in simulation. The half space below the membrane was sampled in the same way, so that coordination numbers include sampling over both.

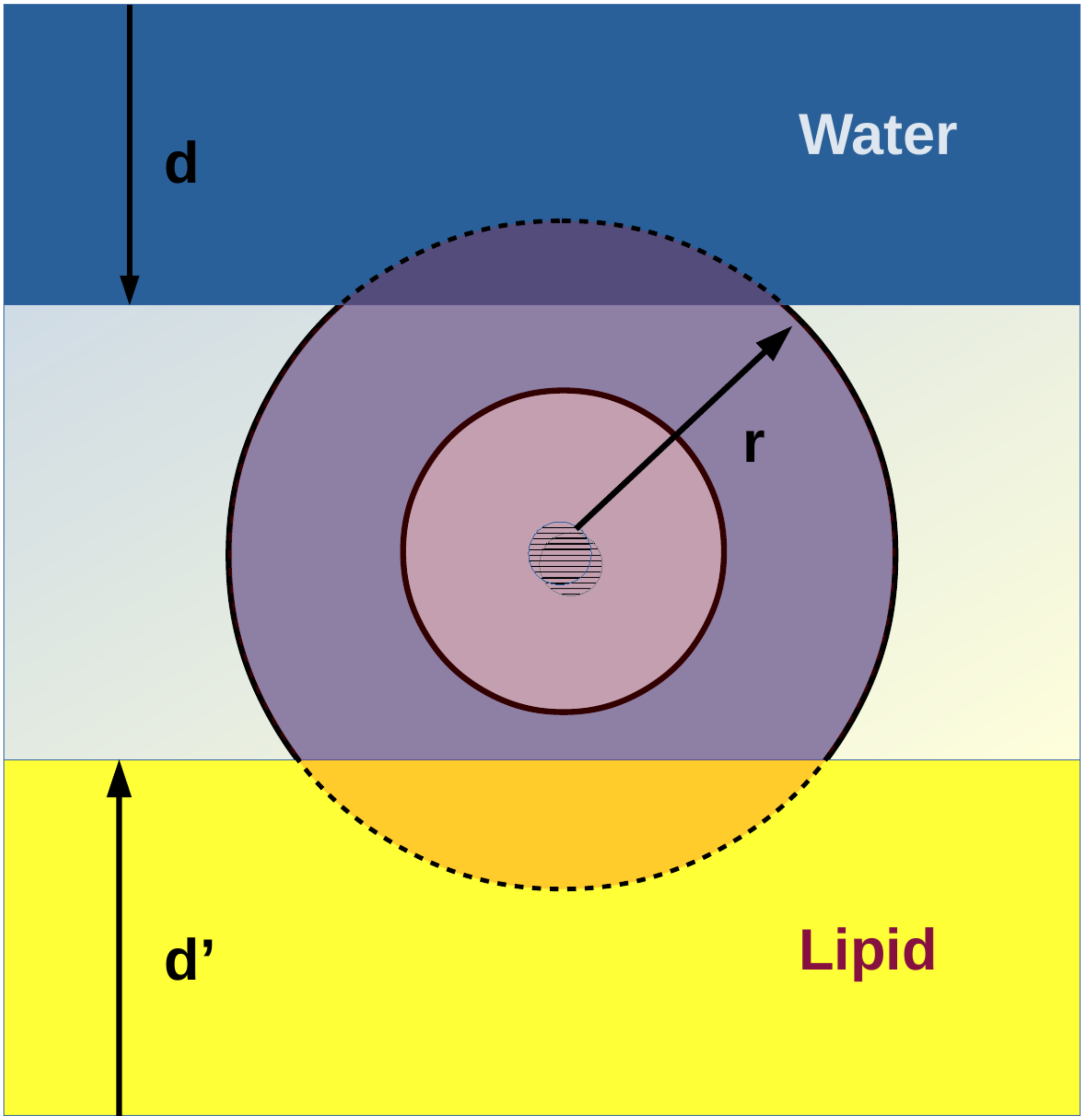


FIG. 3. Schematic for the calculation of the interfacial RDF. At small $r$, the calculation is identical to normal spatial distribution functions. At large $r$, parts of the bulk and the lipid interior are not included, restricting the function to the interface.

#### B. Results for the Interfacial RDF

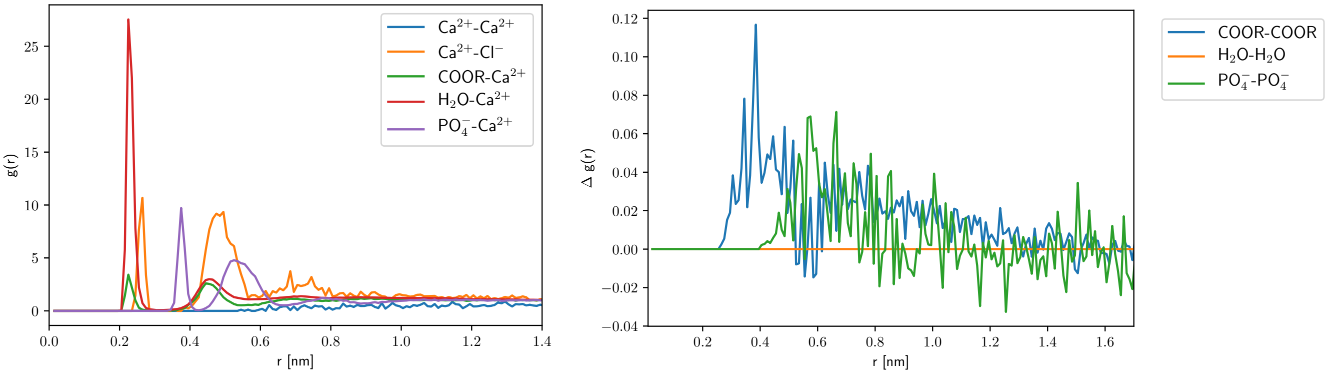


FIG. 4. Interfacial structure of POPC. The coordination numbers are P-$Ca^{2+}$ 0.0 COOR-$Ca^{2+}$ 0.0 At first peak and P-$Ca^{2+}$ 0.1 COOR-$Ca^{2+}$ 0.0 over all shells. Left panel: $Ca^{2+}$ coordination. Right panel: Change in membrane structure upon $Ca^{2+}$ adsorption as given by local RDF change.

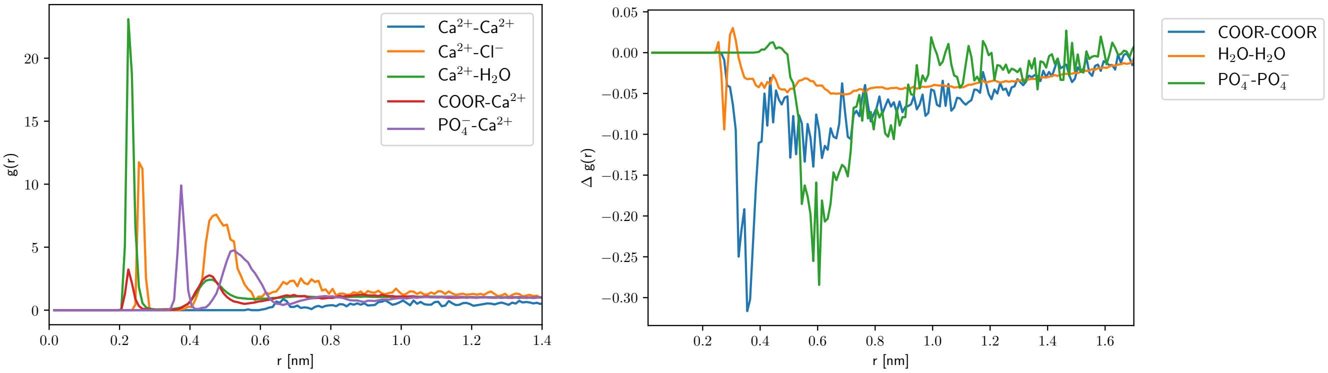


FIG. 5. Interfacial structure of DMPC. The coordination numbers are P-$Ca^{2+}$ 0.0 COOR-$Ca^{2+}$ 0.0 At first peak and P-$Ca^{2+}$ 0.1 COOR-$Ca^{2+}$ 0.0 over all shells. Left panel: $Ca^{2+}$ coordination. Right panel: Change in membrane structure upon $Ca^{2+}$ adsorption as given by local RDF change.

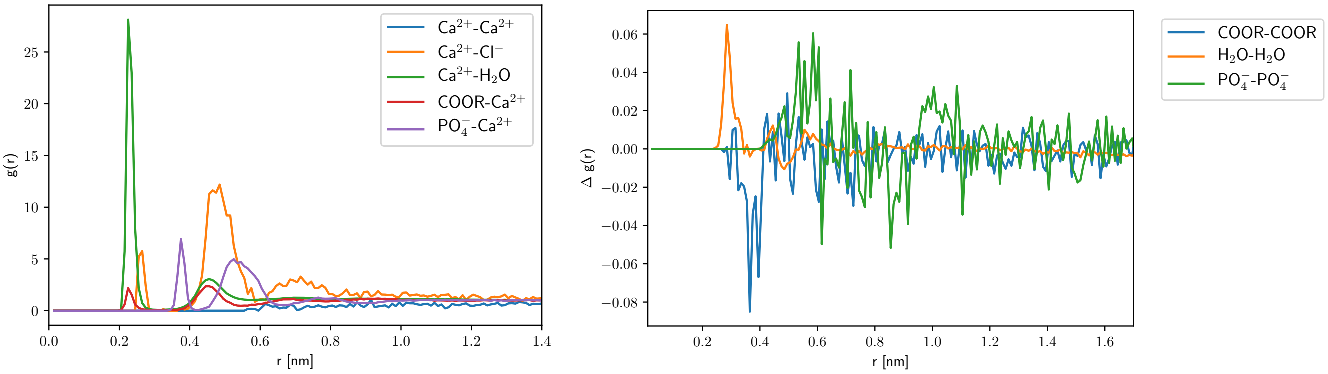


FIG. 6. Interfacial structure of DOPC. The coordination numbers are COOR-$Ca^{2+}$ 0.0 P-$Ca^{2+}$ 0.0 At first peak and COOR-$Ca^{2+}$ 0.0 P-$Ca^{2+}$ 0.1 over all shells. Left panel: $Ca^{2+}$ coordination. Right panel: Change in membrane structure upon $Ca^{2+}$ adsorption as given by local RDF change.

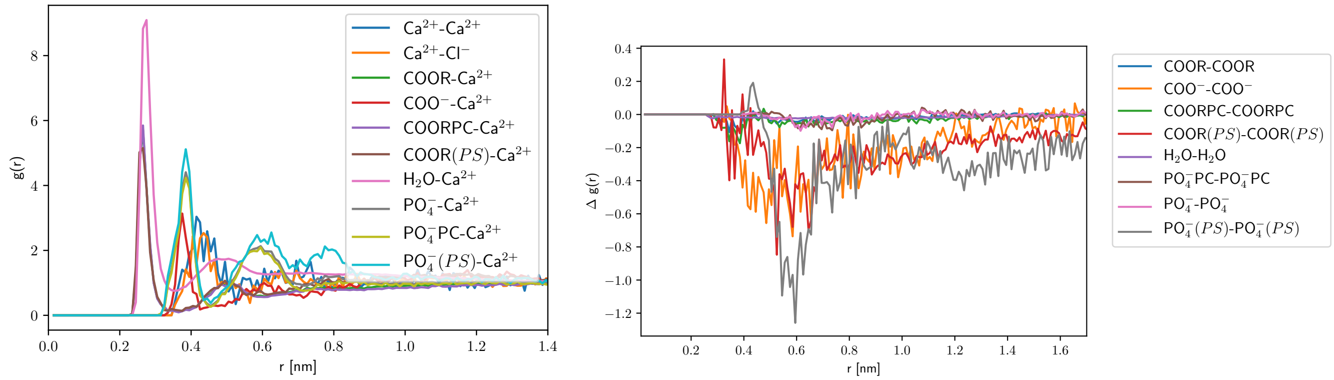


FIG. 7. Interfacial structure of DOPC/DOPS. The coordination numbers are $PO_4^-(PC)$-$Ca^{2+}$ 0.1, COOR($PS$)-$Ca^{2+}$ 0.1. At first peak and $PO_4^-(PS)$-$Ca^{2+}$ 3.6, COOR($PS$)-$Ca^{2+}$ 3.5 over all shells. Left panel: $Ca^{2+}$ coordination. Right panel: Change in membrane structure upon $Ca^{2+}$ adsorption as given by local RDF change.

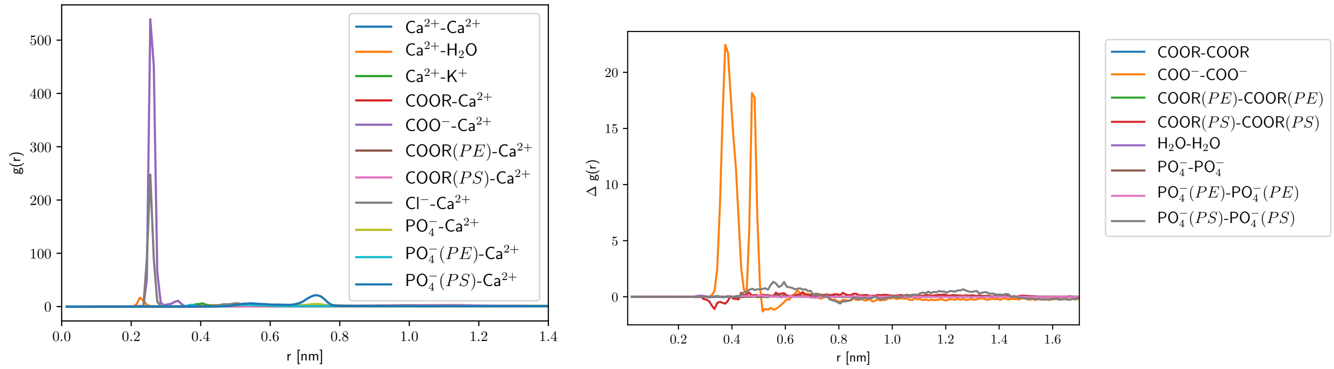


FIG. 8. Interfacial structure of DOPE/DOPS. The coordination numbers are $COO^-$-$Ca^{2+}$ 0.4 $Ca^{2+}$-$K^+$ 0.3 At first peak and $PO_4^-(PS)$-$Ca^{2+}$ 1.0 $COO^-$-$Ca^{2+}$ 0.6 over all shells. Left panel: $Ca^{2+}$ coordination. Right panel: Change in membrane structure upon $Ca^{2+}$ adsorption as given by local RDF change.

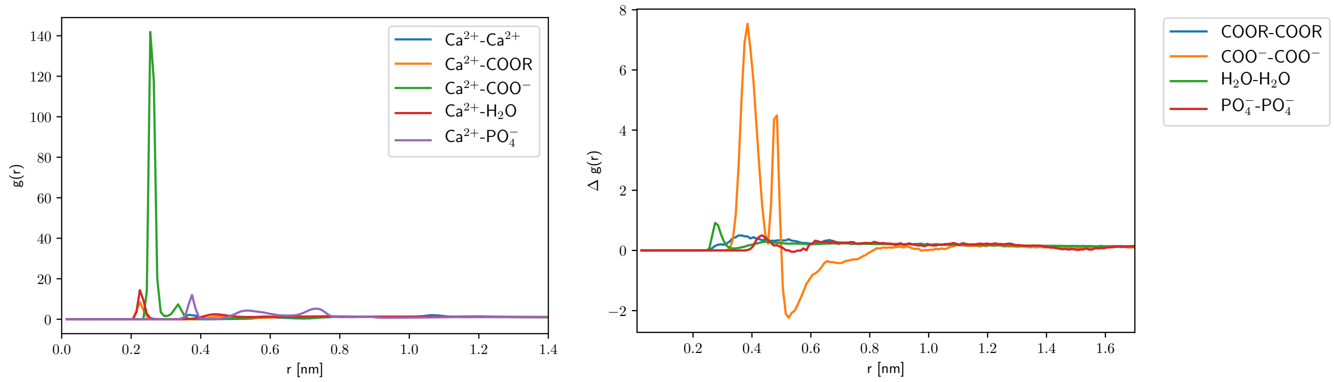


FIG. 9. Interfacial structure of DOPS. The coordination numbers are $Ca^{2+}$-$COO^-$ 0.8 $Ca^{2+}$-COOR 0.6 At first peak and $Ca^{2+}$-$PO_4^-$ 4.5, $Ca^{2+}$-$COO^-$ 2.8 over all shells. Left panel: $Ca^{2+}$ coordination. Right panel: Change in membrane structure upon $Ca^{2+}$ adsorption as given by local RDF change.

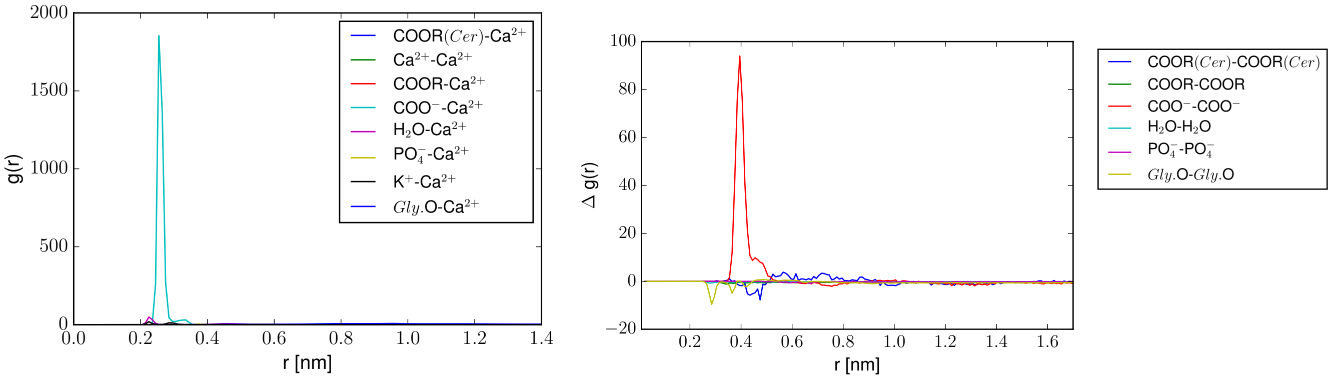


FIG. 10. Interfacial structure of GM1/POPC. The coordination numbers are COOR($PC$)-Ca$^{2+}$ 1.4, COOR($Cer$)-Ca$^{2+}$ 1.0 At first peak and $Gly$.O-Ca$^{2+}$ 5.8, COO$^{-}$-Ca$^{2+}$ 0.7 over all shells. Here $Gly$. refers to all neutral sugar oxygens, and COO$^{-}$ to the sialic acid carboxylate. Left panel: Ca$^{2+}$ coordination. Right panel: Change in membrane structure upon Ca$^{2+}$ adsorption as given by local RDF change.

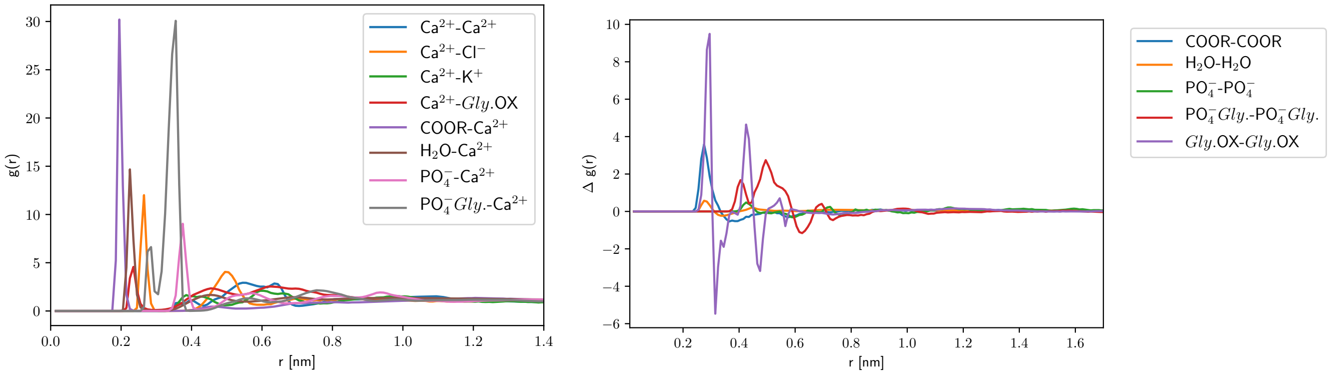


FIG. 11. Interfacial structure of PIP2. The coordination numbers are pip2-ca-P$Gly$.-Ca$^{2+}$ 1.8 P-Ca$^{2+}$ 0.3 At first peak and P$Gly$.-Ca$^{2+}$ 2.5, COOR-Ca$^{2+}$ 2.5 over all shells. Here $Gly$. refers to all neutral sugar oxygens. Left panel: Ca$^{2+}$ coordination. Right panel: Change in membrane structure upon Ca$^{2+}$ adsorption as given by local RDF change.

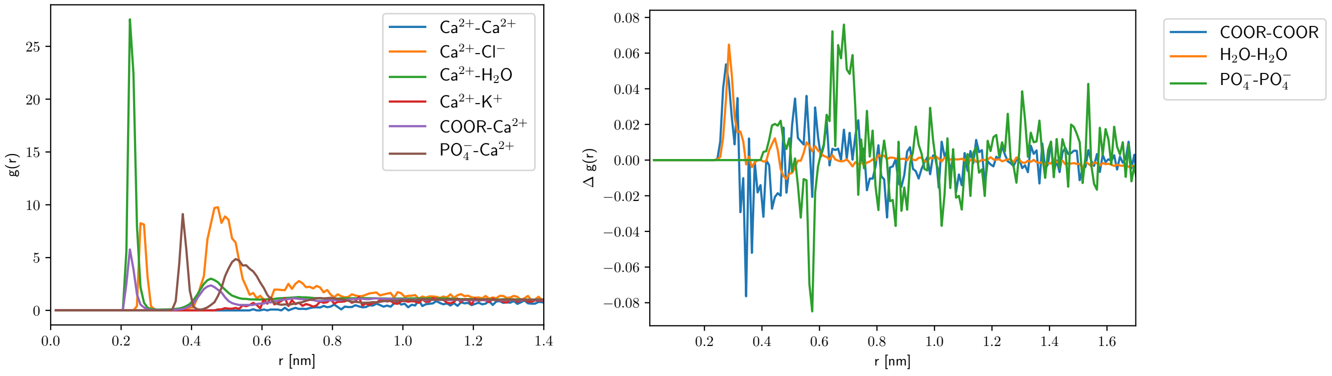


FIG. 12. Interfacial structure of DOPC in presence of KCl. The coordination numbers are Ca$^{2+}$-K$^{+}$ 0.3 COOR-Ca$^{2+}$ 0.0 At first peak and P-Ca$^{2+}$ 0.1 Ca$^{2+}$-K$^{+}$ 0.1 over all shells. Left panel: Ca$^{2+}$ coordination. Right panel: Change in membrane structure upon Ca$^{2+}$ adsorption as given by local RDF change.

TABLE I. System Compositions: Integer Charges

| Name | Lipids | Water | $Ca^{2+}$ | $Cl^-$ | $K^+$ | T[K] |
|---|---|---|---|---|---|---|
| DMPC+$Ca^{2+}$ | DMPC 128 | 4928 | 12 | 23 | | 303 |
| DMPC+$Ca^{2+}$ | DMPC 128 | 4928 | 12 | 23 | | 313 |
| DMPC | DMPC 128 | 4940 | | 11 | 12 | 313 |
| DOPC+$Ca^{2+}$ | DOPC 128 | 4986 | 12 | 24 | | 303 |
| DOPC+KCl+$Ca^{2+}$ | DOPC 128 | 4963 | 12 | 36 | 12 | 303 |
| DOPC+KCl | DOPC 128 | 4999 | | 12 | 12 | 303 |
| DOPC | DOPC 128 | 5023 | | | | 303 |
| DOPC+DOPS+$Ca^{2+}$ | DOPC 112 DOPS 28 | 5443 | 14 | 12 | 12 | 303 |
| DOPC+DOPS | DOPC 112 DOPS 28 | 5443 | | 12 | 40 | 303 |
| DOPE+DOPS+$Ca^{2+}$ | DOPS 28 DOPE 112 | 5153 | 14 | 12 | 12 | 303 |
| DOPE+DOPS | DOPS 28 DOPE 112 | 5153 | | 12 | 40 | 303 |
| DOPS+$Ca^{2+}$ | DOPS 128 | 5192 | 64 | | | 303 |
| DOPS | DOPS 128 | 5192 | | | 128 | 303 |
| POPC+GM1 | GM1 26 POPC 104 | 9978 | | | 26 | 313 |
| POPC+GM1+$Ca^{2+}$ | GM1 26 POPC 104 | 9978 | 13 | | | 313 |
| PIP2+$Ca^{2+}$ | SAPI2 128 | 6600 | 256 | 16 | 16 | 313 |
| PIP2 | SAPI2 128 | 6600 | | 16 | 528 | 313 |
| POPC+$Ca^{2+}$ | POPC 128 | 4863 | 12 | 24 | | 313 |
| POPC | POPC 128 | 4863 | | | | 313 |

TABLE II. System Compositions: Scaled Charges

| Name | Lipids | Water | $Ca^{2+}$ | $Cl^-$ | $Na^+$ | T[K] |
|---|---|---|---|---|---|---|
| DMPC+$Ca^{2+}$ | DMPC 128 | 4928 | 12 | 23 | | 303 |
| DOPC+$Ca^{2+}$ | DOPC 128 | 4986 | 12 | 24 | | 303 |
| DOPC+KCl+$Ca^{2+}$ | DOPC 127 | 4963 | 12 | 36 | 12 | 303 |
| DOPC+DOPS+$Ca^{2+}$ | DOPS 28 DOPE 112 | 5153 | 14 | 2 | 12 | 303 |
| DOPE+DOPS+$Ca^{2+}$ | DOPS 28 DOPE 112 | 5153 | 14 | 2 | 12 | 303 |

### IV. ADDITIONAL SIMULATION DETAILS

All systems were pre-equilibrated for at least 200 ns and then sampled for an additional 200 ns, the PIP2 system was pre-equilibrated for 400 ns, until the APL stabilized. We used CHARMM-GUI[22] to create the initial setups, a timestep of 0.5 fs and a Nosé-Hoover[23, 24] thermostat with a coupling constant of 1.0 ps, as well as a semiisotropic Parrinello-Rahman[25] barostat, with a timeconstant of 5 ps. The PME[26] cutoff was 1.1 nm as was the Lennard-Jones potential cutoff. No potential smoothing or dispersion corrections were used, as these are not supported by the lateral pressure codes. All sampling was done using GROMACS 5.[18] For initial equilibration, we sometimes used OpenMM[27], with identical cutoffs. Hydrogen bonds were constrained using LINCS[17]. We used the TIP3P[28] forcefield for water and the CHARMM36[29] forcefield for lipids.

### V. SYSTEM COMPOSITIONS

### VI. GEL PHASE

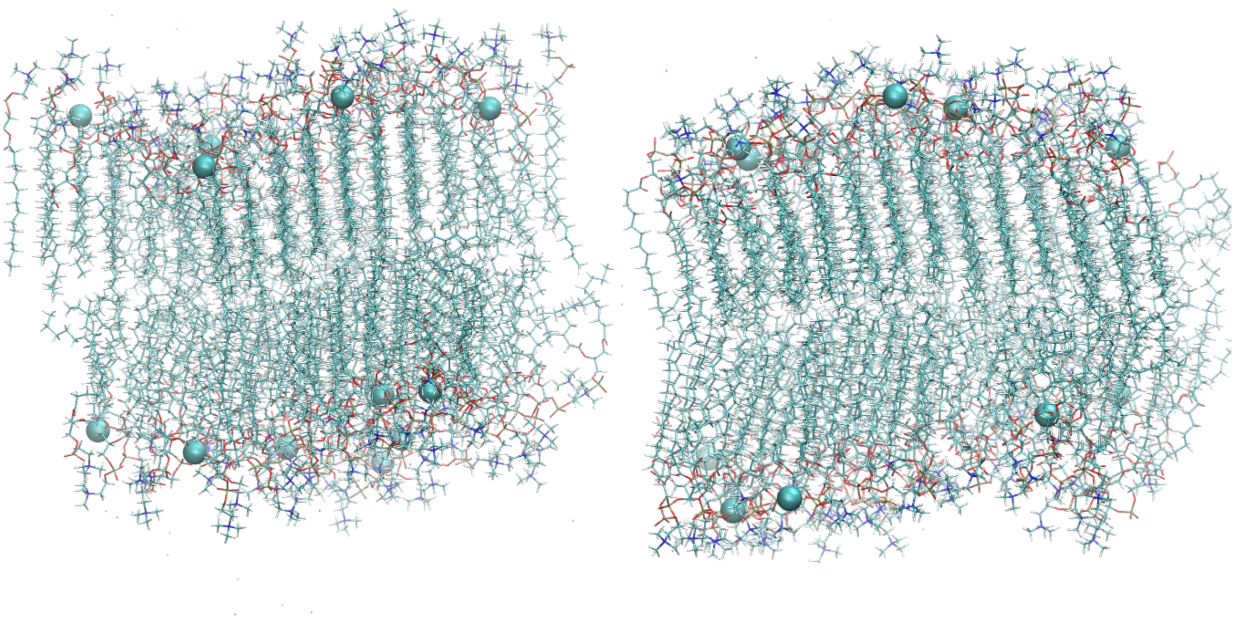

FIG. 13. Gel phase of DMPC at 303 K in presence of $Ca^{2+}$. Left panel: Standard CHARMM forcefield and ions. Right panel: Simulatoin with scaled charges. In the absence of calcium ions, the membrane was fluid, the melting temperature for pure DMPC is ca. 297 K.

### VII. INTERFACE AND COORDINATION

In Table III, we give additional details on the interface and its coordination structure. The Gibbs-Luzzati plane was obtained from the water density by finding the surface of zero excess for water using

$$\int_{-\infty}^{z_{GDS}} c_{H_2O}(z)dz + \int_{z_{GDS}}^{\infty} c_0 - c_{H_2O}(z) = \Gamma_{H_2O} = 0 \tag{35}$$

where $c_0$ is the water concentration in the middle between the two bilayers, and $z$ is the coordinate normal to the membrane surface. $z_{GDS}$ was determined as the $z$ coordinate where the water excess $\Gamma_{H_2O}$ was zero. In our boxes the infinities were replaced by the box dimension and the midplane respectively. ReSIS thicknesss was computed using the $z$ positions of the membrane lipids as computed from the lipid definitions used for the ReSIS surfaces[21]. The peak positions of the PO4 headgroup were estimated from the maximum number density, their width was computed by taking two standard deviations of their density distribution. We used symmetrized density distributions for this purpose. We computed the hydration force decay length $\xi_0$ from the water orientation $\eta(x) = \langle \mathbf{p} \cdot x \rangle$, by fitting the equation

$$\eta(x) = \eta_0 \frac{\sinh x\xi_0^{-1}}{\sinh d/2\xi_0^{-1}} \tag{36}$$

according following Ref. [30], inspired by the work of Marčelja and Radić[31]. The fitting zone was limited by a water excess of 500. Results are given in Table V.

TABLE III. Description of $PO_4$ headgroup position and distribution, as well as Gibbs-Luzzatti plane and a ReSIS estimate of the pivotal plane.

| Name | Peak [nm] | Width [nm] | Height [N/nm$^3$] | Luzzatti [nm] | ReSIS [nm] |
|---|---|---|---|---|---|
| DMPC+$Ca^{2+}$ | 1.83 | 0.53 | 2.74 | 1.98 | 1.16 |
| DMPC | 1.76 | 0.59 | 2.38 | 1.86 | 1.16 |
| DOPC+KCl | 1.91 | 0.49 | 2.42 | 1.84 | 1.29 |
| DOPC | 1.79 | 0.46 | 2.44 | 1.96 | 1.28 |
| DOPC+DOPS+$Ca^{2+}$ | 2.02 | 0.49 | 2.63 | 2.09 | 1.34 |
| DOPC+DOPS | 1.89 | 0.46 | 2.55 | 1.94 | 1.31 |
| DOPE+DOPS+$Ca^{2+}$ | 2.06 | 0.49 | 2.70 | 2.10 | 1.30 |
| DOPE+DOPS | 1.98 | 0.45 | 2.76 | 2.05 | 1.27 |
| DOPS+$Ca^{2+}$ | 2.03 | 0.47 | 2.83 | 2.13 | 1.41 |
| DOPS | 1.94 | 0.45 | 2.79 | 1.99 | 1.36 |
| POPC+GM1 | 1.95 | 0.54 | 1.93 | 2.41 | 1.31 |
| POPC+GM1+$Ca^{2+}$ | 1.81 | 0.51 | 1.94 | 2.23 | 1.32 |
| POPC+GM1 | 1.98 | 0.49 | 2.10 | 2.26 | 1.32 |
| PIP2+$Ca^{2+}$ | 1.87 | 0.62 | 1.95 | 2.13 | 1.54 |
| PIP2 | 1.82 | 0.45 | 2.47 | 2.08 | 1.49 |
| POPC+$Ca^{2+}$ | 1.75 | 0.56 | 2.22 | 2.03 | 1.28 |
| POPC | 1.87 | 0.50 | 2.41 | 1.93 | 1.27 |

TABLE IV. $Ca^{2+}$ binding characterized by the number density peaks (symmetrized)

| Name | Peak [nm] | Width [nm] | Height [N/nm$^3$] |
|---|---|---|---|
| DMPC+$Ca^{2+}$ | 1.53 | 0.42 | 0.31 |
| DOPC+DOPS+$Ca^{2+}$ | 1.87 | 0.67 | 0.24 |
| DOPE+DOPS+$Ca^{2+}$ | 1.86 | 0.44 | 0.30 |
| DOPS+$Ca^{2+}$ | 2.05 | 0.53 | 1.29 |
| POPC+GM1+$Ca^{2+}$ | 2.45 | 1.49 | 0.11 |
| PIP2+$Ca^{2+}$ | 2.26 | 0.78 | 3.21 |
| POPC+$Ca^{2+}$ | 1.25 | 1.24 | 0.09 |

TABLE V. Ideal Osmotic pressures, computed from concentrations inside a window around the middle of the water phase. Marčelja decay length $\xi_0$

| Name | cRT [bar] | [$K^+$] mol/l | [$Cl^-$] mol/l | [$Ca^{2+}$] mol/l | Marčelja $\xi_0$ [nm] |
|---|---|---|---|---|---|
| DMPC+$Ca^{2+}$ | 2.790 | 0.000 | 0.067 | 0.000 | n/a |
| DMPC | 3.739 | 0.037 | 0.053 | 0.000 | 1.130 |
| DOPC | 0.000 | 0.000 | 0.000 | 0.000 | 1.361 |
| DOPC+KCl | 4.026 | 0.040 | 0.056 | 0.000 | 1.327 |
| DOPC+DOPS+$Ca^{2+}$ | 3.555 | 0.037 | 0.048 | 0.000 | 3.481 |
| DOPC+DOPS | 5.979 | 0.087 | 0.055 | 0.000 | 1.640 |
| DOPE+DOPS+$Ca^{2+}$ | 3.395 | 0.039 | 0.042 | 0.000 | 1.295 |
| DOPE+DOPS | 6.140 | 0.092 | 0.055 | 0.000 | 4.876 |
| DOPS+$Ca^{2+}$ | 0.000 | 0.000 | 0.000 | 0.000 | 1.320 |
| DOPS | 3.761 | 0.090 | 0.000 | 0.000 | 7.972 |
| POPC+GM1 | 2.046 | 0.049 | 0.000 | 0.000 | n/a |
| POPC+GM1+$Ca^{2+}$ | 3.964 | 0.000 | 0.000 | 0.095 | n/a |
| PIP2+$Ca^{2+}$ | 1.500 | 0.005 | 0.031 | 0.000 | 0.545 |
| PIP2 | 8.185 | 0.123 | 0.073 | 0.000 | 7.361 |
| POPC+$Ca^{2+}$ | 5.725 | 0.000 | 0.103 | 0.034 | 0.845 |
| POPC | 0.000 | 0.000 | 0.000 | 0.000 | 5.911 |

TABLE VI. Computed Monolayer Bending Moments and Bilayer Gaussian Moduli from Harasima Stress Tensor

| Name | $\kappa_m J_s$ [kT]/nm$^{-1}$ | $\bar{\kappa_b}$ [kT] |
|---|---|---|
| POPC | -0.01 | -7.98 |
| DMPC | -1.15 | -0.49 |
| DOPC | -0.35 | -3.45 |
| DOPC+DOPS | -0.84 | -2.42 |
| DOPE+DOPS | -3.49 | 13.93 |
| DOPS | -1.58 | -4.92 |
| POPC+GM1 | 1.59 | -34.49 |
| PIP2 | -1.86 | -2.81 |
| POPC+$Ca^{2+}$ | -0.08 | -6.60 |
| DMPC+$Ca^{2+}$ | -1.68 | 6.37 |
| DOPC+$Ca^{2+}$ | -0.01 | -6.12 |
| DOPC+DOPS+$Ca^{2+}$ | -1.59 | 1.44 |
| DOPE+DOPS+$Ca^{2+}$ | -4.57 | 21.45 |
| DOPS+$Ca^{2+}$ | -4.43 | 17.83 |
| POPC+GM1+$Ca^{2+}$ | -0.11 | -11.04 |
| PIP2+$Ca^{2+}$ | -8.89 | 56.20 |

TABLE VII. Elastic Constants and Areas per lipid

| Name | $\kappa$ [kT] | $\kappa_\theta^b$ [kT/nm$^2$] | $A_l$ [$\AA^2$] | STDEV [$\AA^2$] | ReSIS Pivot [nm] |
|---|---|---|---|---|---|
| POPC | 12.93 | 15.20 | 65.80 | 1.41 | 1.27 |
| POPC+$Ca^{2+}$ | 13.89 | 15.42 | 64.83 | 1.47 | 1.28 |
| DMPC | 17.00 | 16.37 | 61.09 | 1.30 | 1.16 |
| DMPC+$Ca^{2+}$ | 17.80 | 20.61 | 61.14 | 1.40 | 1.16 |
| DOPC | 11.50 | 14.73 | 67.90 | 1.30 | 1.28 |
| DOPC+$Ca^{2+}$ | 11.88 | 15.46 | 67.57 | 1.39 | 1.29 |
| DOPC+DOPS | 12.75 | 15.16 | 67.08 | 1.06 | 1.31 |
| DOPC+DOPS+$Ca^{2+}$ | 11.86 | 16.26 | 65.98 | 1.18 | 1.30 |
| DOPE+DOPS | 16.41 | 16.30 | 61.35 | 1.07 | 1.27 |
| DOPE+DOPS+$Ca^{2+}$ | 16.67 | 24.59 | 60.98 | 1.13 | 1.28 |
| DOPS | 13.89 | 15.56 | 64.25 | 1.22 | 1.36 |
| DOPS+$Ca^{2+}$ | 17.63 | 25.53 | 60.74 | 1.04 | 1.41 |
| POPC+GM1 | 16.92 | 21.74 | 61.38 | 1.45 | 1.31 |
| POPC+GM1+$Ca^{2+}$ | 16.98 | 22.79 | 60.94 | 1.10 | 1.32 |
| PIP2 | 12.96 | 13.86 | 72.13 | 1.30 | 1.49 |
| PIP2+$Ca^{2+}$ | 14.70 | 19.95 | 68.56 | 0.34 | 1.54 |

[1] Y. Kozlovsky and M. M. Kozlov, Biophys. J. **82**, 882 (2002).
[2] M. Hamm and M. Kozlov, Eur. Phys. J. E **3**, 323 (2000).
[3] Y. Kozlovsky, A. Efrat, D. A. Siegel, and M. M. Kozlov, Biophys. J. **87**, 2508 (2004).
[4] P. Virtanen, R. Gommers, T. E. Oliphant, M. Haberland, T. Reddy, D. Cournapeau, E. Burovski, P. Peterson, W. Weckesser, J. Bright, et al., arXiv e-prints arXiv:1907.10121 (2019), 1907.10121.
[5] J. H. Irving and J. G. Kirkwood, J. Chem. Phys. **18**, 817 (1950).
[6] H.-J. Kreuzer, *Nonequilibrium thermodynamics and its statistical foundations.* (Clarendon Press, Oxford, 1981).
[7] P. Schofield, J. R. Henderson, and J. S. Rowlinson, Proc. R. Soc. A **379**, 231 (1982).
[8] W. Noll, J. Rational Mech. Anal. **4**, 627 (1955).
[9] A. Harasima, Adv. Chem. Phys. **1**, 203 (1958).
[10] J. Sonne, F. Y. Hansen, and G. H. Peters, J. Chem. Phys. **122**, 124903 (2005).
[11] M. Sega, B. Fábián, and P. Jedlovszky, J. Chem. Theory Comput. **12**, 4509 (2016).
[12] R. Goetz and R. Lipowsky, J. Chem. Phys. **108**, 7397 (1998).
[13] J. M. Vanegas, A. Torres-Sánchez, and M. Arroyo, J. Chem. Theory Comput. **10**, 691 (2014).
[14] J. Rigelesaiyin, A. Diaz, W. Li, L. Xiong, and Y. Chen, Proc. R. Soc. A **474**, 20180155 (2018).
[15] H.-S. Lee, Bull. Korean Chem. Soc. **33**, 3039 (2012).
[16] S. Miyamoto and P. A. Kollman, J. Comp. Chem. **13**, 952 (1992), https://onlinelibrary.wiley.com/doi/pdf/10.1002/jcc.540130805, URL `https://onlinelibrary.wiley.com/doi/abs/10.1002/jcc.540130805`.
[17] B. Hess, H. Bekker, H. J. C. Berendsen, and J. G. E. M. Fraaije, J. Comp. Chem. **18**, 1463 (1997), ISSN 1096-987X.
[18] M. J. Abraham, T. Murtola, R. Schulz, S. Páll, J. C. Smith, B. Hess, and E. Lindahl, SoftwareX **1–2**, 19 (2015), ISSN 2352-7110.
[19] A. Ben-Naim, *Molecular Theory of Solutions* (Oxford University Press, New York, 2006).
[20] A. D. Polyanin and A. V. Manzhirov, *Mathematical Handbook for Engineers and Scientists* (Taylor & Francis, 2006), p. 69.
[21] C. Allolio, A. Haluts, and D. Harries, Chem. Phys. **514**, 31 (2018).
[22] S. Jo, T. Kim, V. G. Iyer, and W. Im, J. Comp. Chem. **29**, 1859 (2008), ISSN 1096-987X.
[23] S. Nosé, Mol. Phys. **5**, 255 (1984).
[24] W. G. Hoover, Phys. Rev. A **31**, 1695 (1985).
[25] M. Parrinello and A. Rahman, Phys. Rev. Lett. **45**, 1196 (1980).
[26] U. Essman, L. Perela, M. L. Berkowitz, T. Darden, H. Lee, and L. G. Pedersen, J. Chem. Phys. **103**, 8577 (1995).
[27] P. Eastman, J. Swails, J. D. Chodera, R. T. McGibbon, Y. Zhao, K. A. Beauchamp, L.-P. Wang, A. C. Simmonett, M. P. Harrigan, C. D. Stern, et al., PLOS Comput. Biol. **13**, 1 (2017), URL `https://doi.org/10.1371/journal.pcbi.1005659`.
[28] W. L. Jorgensen, J. Chandrasekhar, J. D. Madura, R. W. Impey, and M. L. Klein, J. Chem. Phys. **79**, 926 (1983).
[29] R. B. Best, X. Zhu, J. Shim, P. E. M. Lopes, J. Mittal, M. Feig, and J. Alexander D. MacKerell, J. Chem. Theory Comput. **8**, 3257 (2012).
[30] M. Kanduč and R. R. Netz, Proc. Natl. Acad. Sci. USA **112**, 12338 (2015).
[31] N. R. S Marčelja, Chem. Phys. Lett. **42**, 129 (1976).
